# Shape2Fate: a morphology-aware deep learning framework for tracking endocytic and exocytic carriers at nanoscale

**DOI:** 10.64898/2026.03.29.715120

**Authors:** Adam Harmanec, Alexander D. Dagg, Jan Kamenicky, Tomas Kerepecky, Yelyzaveta Makieieva, Conceição Pereira, Nicholas Bright, Dilip Menon, David C. Gershlick, Nadezda Vaskovicova, Tiffany Lai, Daniel Fazakerley, Lothar Schermelleh, Filip Sroubek, Zuzana Kadlecova

## Abstract

Plasma membrane homeostasis requires balanced exocytosis and endocytosis, yet their coordination at the single-event level in non-neuronal cells is unresolved. We present Shape2Fate, a morphology-aware deep-learning pipeline that detects, tracks, and classifies individual exocytic and endocytic carriers in live-cell total internal reflection fluorescence structured illumination microscopy (TIRF-SIM) movies at ∼100 nm resolution. Trained on synthetic data and exploiting carrier shape evolution rather than fluorescence intensity, Shape2Fate achieves expert-level tracking and outcome classification across diverse cell types, imaging conditions, and microscope platforms. Applying Shape2Fate to constitutive secretion and insulin-stimulated GLUT4 exocytosis in adipocytes, we uncover two opposing exo-endocytic coupling architectures: exocytic fusion locally nucleates de novo clathrin-coated pits, whereas GLUT4 vesicles target pre-existing pits for rapid cargo capture. These findings establish that the spatial rules governing exo-endocytic coordination are not universal but are pathway-specific. Shape2Fate is openly available, enabling direct event-level mechanistic dissection of exo-endocytic coordination across pathways in living cells.

## Introduction

Plasma membrane (PM) homeostasis depends on the dynamic balance between exocytosis and endocytosis [1]. At neuronal synapses, this coupling is experimentally tractable and has therefore been dissected in depth. Vesicle turnover is rapid and repetitive, confined to stable micron-scale release zones, and accessible to complementary electrophysiological readouts [2–4]. By contrast, in most non-neuronal cells, secretory delivery, recycling and clathrin-mediated endocytosis (CME) are dispersed across large, heterogeneous membrane areas. Whether these pathways are coordinated locally and globally, and if so, with what spatial and temporal hierarchy, remains poorly defined. To resolve this, it is necessary to detect large numbers of individual exo- and endocytic events, localise them with sub-diffraction precision, link them into trajectories and show statistically that their co-ordination in space and time exceeds chance expectations. Yet automated single-event tracking in live-cell fluorescence movies is hampered by crowding, heterogeneous appearances of membrane trafficking carriers, motion blur and rapid shape transitions that confound object identity and can obscure productive cargo delivery.

High-contrast, membrane-proximal imaging is essential for studying PM homeostasis. Although TIRF microscopy provides this selectivity, its ∼200-300 nm lateral resolution obscures carrier morphology and merges nearby events. As a result, intensity-based TIRF tracking often cannot resolve individual carrier trajectories or classify outcomes (productive vs failed cargo delivery). The challenge is acute because membrane trafficking processes produce a spectrum of outcomes that differ morphologically but overlap in intensity. In CME, CCPs can abort via premature disassembly [5]. Exocytic carriers, on the other hand, may fail to dock or complete fusion at the PM and thus never deliver their cargo [6, 7]. Resolving productive from failed events at the single-event level is therefore essential: the fraction of attempts that succeed determines the net membrane flux, and only by identifying individual productive events can we map their spatial and temporal organisation relative to other cellular processes. However, productive and abortive events have broadly overlapping lifetimes and peak intensities [5, 8]. Moreover, measured fluorescence intensity depends on probe labelling, photobleaching rates, and acquisition settings. Individual intensity–time traces cannot reliably distinguish event outcomes. Population-level intensity or lifetime thresholds do not resolve this ambiguity either, as they are calibrated to specific acquisition conditions and do not transfer reliably across different experimental systems and microscope platforms [8]. Together, these limitations motivate event-by-event analyses that require an orthogonal, transferable productivity criterion applicable across pathways and experimental systems, while preserving single-carrier heterogeneity. TIRF combined with 2D structured illumination microscopy (TIRF-SIM) has emerged as a solution that meets the required spatiotemporal resolution [9–11].

TIRF-SIM offers ∼100 nm spatial resolution while retaining high temporal resolution and live-cell imaging compatibility, thus enabling direct visualization of endocytic pit maturation [12, 13] and, in principle, the distinct morphological transitions of exocytic carriers throughout their lifetime. This resolution gain comes at the cost of computationally reconstructing each frame from multiple phase- and orientation-shifted raw images [11], rendering the data susceptible to reconstruction artefacts [14–16]. Such artefacts arise from vesicle motion during raw-frame acquisition, low signal-to-noise ratios (SNR) in live cells, and imperfect illumination-pattern estimation. These SIM reconstruction artefacts add to the intrinsic challenges of carrier complex morphologies and their overlap, complicating detection and trajectory linking. As a result, methods such as thresholding or Gaussian fitting [17–19], already limited in standard TIRF, perform even less reliably on artefact-prone TIRF-SIM data [12, 20].

Overcoming these specific challenges requires more sophisticated multi-object tracking (MOT) methods for live-cell microscopy [21–23]. While MOT methods have been introduced for single molecule localization microscopy [24], a dedicated MOT framework for complex objects observed in super-resolution microscopy has not yet been established. The prevailing MOT strategy is “tracking-by-detection”: objects are detected independently in each frame and then linked into trajectories over time. Typically, detections are either associated frame-to-frame [17] or by formulating the linking problem as a global optimization [22, 25]. Linking often fails during rapid shape transitions or in crowded scenes with overlapping objects [20, 21]. Yet these same morphological transitions are biologically informative. Rather than changing arbitrarily, trafficking carriers follow reproducible morphological trajectories that reflect their progression toward distinct functional outcomes, motivating a morphology-aware MOT solution. Existing trackers in biological imaging of sub-cellular structures typically treat all detections uniformly and rely on motion models and spatial proximity for association [17, 18, 26]. In natural-image MOT (vehicle or pedestrian tracking for example) association is often strengthened using learned appearance embeddings [27]. Related feature-learning strategies have influenced recent whole-cell tracking frameworks, including EmbedTrack [28] and transformer-based Trackastra [23]. However, these methods operate at the whole-cell scale and are not optimized for the nanoscale variability and subsecond shape changes of subcellular objects in super-resolution microscopy. Nor do they explicitly exploit how object shape evolves over an entire trajectory, which is the key feature that can distinguish productive from failed cargo delivery.

Deep-learning-based detectors [29, 30] and trackers [23, 28] can better adapt to morphology and its transitions, but typically require extensive labelled training data and careful tuning, with model parameters often needing adjustment for each new imaging condition or experimental setup [31]. To avoid time- and resource-intensive manual annotation of microscopy time series, an alternative is to generate synthetic data that approximate the structure of the biological object and simulate the image formation process of microscopes. Such synthetic data can be produced either through mathematical modelling [32, 33] or by using generative approaches such as GANs [34, 35] or, more recently, diffusion models [36].

Motivated by these challenges, we developed Shape2Fate, a fully automated deep-learning pipeline for TIRF-SIM imaging, trained on synthetic data. Shape2Fate integrates shape classification into detection and linking, enabling identity-preserving trajectories with per-event productivity labels. Using Shape2Fate, thoroughly validated and benchmarked against human annotations and existing TIRF tracking pipelines, we mapped exo-endocytic coordination across local and global scales in two mechanistically distinct systems. The first is a model of constitutive secretion, engineered using the RUSH (Retention Using Selective Hooks) system [37], in which biotin addition triggers synchronised release of secretory carriers from the ER [38]. Although RUSH is widely used to study constitutive trafficking, the impact of a cell-wide wave of secretion on plasma membrane homeostasis and endocytic dynamics has not been examined. As a second model, we interrogated insulin-stimulated Glucose Transporter Type 4 (GLUT4) exocytosis in adipocytes, a hormone-regulated pathway in which GLUT4 storage vesicles (GSVs) fuse with the PM to regulate glucose uptake. Impaired GLUT4 trafficking in muscle and adipose tissue underlies insulin resistance, a major risk factor for type 2 diabetes and other metabolic diseases [39]. Shape2Fate reveals that these two systems employ diametrically opposed coupling architectures: in synchronized constitutive secretion, productive exocytic fusion is followed by local de novo CCP formation, whereas in adipocytes insulin-stimulated GSV fusion targets pre-existing CCPs. These findings establish that the spatial rules linking exocytosis to endocytosis are not universal but are instead tuned to the specific demands of each trafficking pathway.

## Results

### Shape2Fate Pipeline Overview

Shape2Fate (Fig. 1A) is an end-to-end pipeline that processes at each time point the nine raw structured-illumination frames (3 angles × 3 phases) together with microscope parameters. It outputs identity-preserving carrier trajectories annotated with per-frame shape labels and a cargo-delivery outcome for each event. The pipeline comprises four modules: (1) SIM reconstruction, (2) object detection with shape classification, (3) global trajectory linking, and (4) quantitative analysis of event outcomes. Each module is engineered for robustness across varying SNRs, cell types, labelling strategies, reconstruction artefacts and microscope configurations, enabling consistent performance across diverse datasets (Fig. 1B, C).

**Figure 1.**
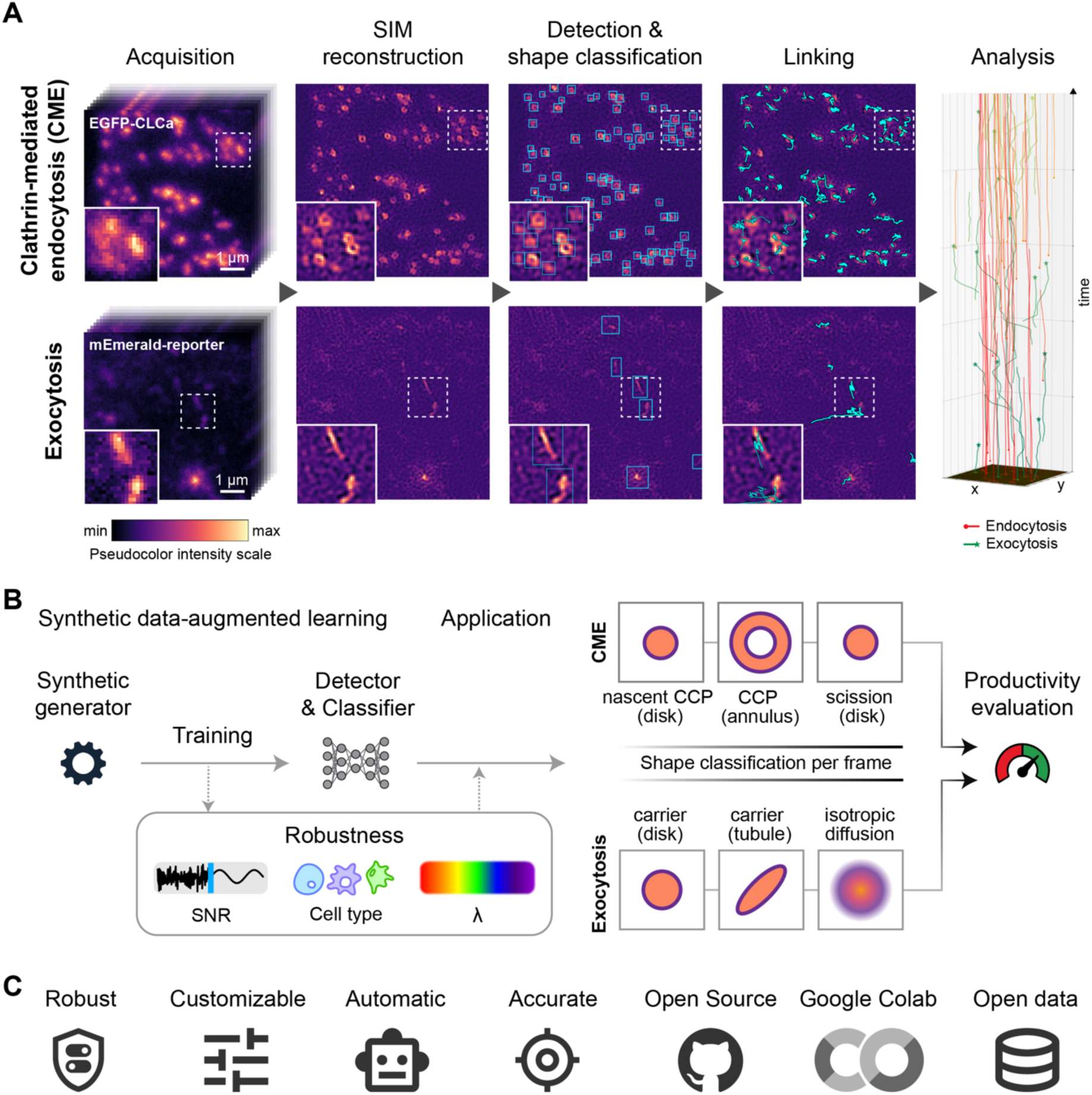
Shape2Fate: a morphology-aware framework that links carrier shape evolution to cargo-delivery outcome. **(A)** TIRF-SIM movies are reconstructed; objects are detected and shape-classified by a neural network; tracks are formed through global linking; and the output is fed into a built-in module for quantitative analysis. Scale bar: 1 µm. **(B)** Training and robustness. A physics-based synthetic generator spans SNR, cell types, fluorophores, and microscope parameters; applied to real data, the model yields simultaneous detection and shape classification to quantify event productivity. **(C)** Design principles and accessibility. Shape2Fate is robust across imaging conditions and object classes, customizable to new cellular processes via the synthetic generator, fully automatic, and achieves human-level tracking accuracy. All code, pre-trained models, Google Colab notebooks, datasets, and documentation are openly available at https://shape2fate.utia.cas.cz/.

At each time point, a super-resolution SIM image is reconstructed using a generalized Wiener filter [40]. To mitigate potential reconstruction artefacts arising from noise, object motion during SIM acquisition, or imperfect illumination-parameter estimates, we address these through three complementary strategies. First, to improve illumination-parameter estimation, we apply a noise-robust procedure [41] to raw SIM frames that have been deconvolved using the Richardson–Lucy algorithm [42]. The SIM reconstruction itself and the entire downstream analysis, however, is performed on the original raw SIM frames. Second, to improve robustness to interframe motion and noise, we train on artefact-containing data (Supplementary Fig. 1). Third, we use linking to correct residual detection errors.

In the reconstructed images, vesicles are localized by a deep neural network trained entirely on realistic synthetic SIM data that emulate object morphology, imaging physics, and artefacts (Fig. 1B). We generated the synthetic dataset by modelling the morphological range observed in our TIRF-SIM and SIM experiments [13, 38, 43]. For CCPs, structures spanned from near-diffraction-limited compact disks to annular profiles with peak-to-peak diameters of ∼120–140 nm in RPE-1 cells. For exocytic carriers we modelled pleomorphic tubules ∼0.5–2 µm in length and up to ∼200 nm in cross-section, together with isotropic signal bursts representing fusion events. We also drew on other published microscopy and electron microscopy (EM) images of similar structures to guide the expected size and shape [12, 44–47]. These synthetic structures were then used to simulate the full TIRF-SIM image-formation process (see Methods). Importantly, we did not impose any specific biochemical or biophysical assembly/disassembly model on these synthetic structures. This data-driven approach focuses solely on matching the morphological patterns in real TIRF-SIM images, following the general principle that synthetic microscopy data can effectively substitute for annotated experimental images in training deep-learning models [36]. Rather than predicting a binary mask, the network outputs a normalized interior-distance map [48, 49], which encodes both object size and morphology. In this representation, each pixel inside an object is assigned a value equal to its shortest normalized distance to the boundary. This enables simultaneous detection and shape inference, even in dense clusters or intermingling carriers. From each detected endocytic or exocytic carrier, we derive a single scalar shape descriptor, termed the Shape Index (SI), which feeds directly into the linking and downstream analysis. This step is readily extensible by updating the synthetic generator, enabling an effectively unlimited supply of training examples so long as the targets fall within the imaging and morphology regime of membrane-trafficking carriers (Fig. 1B).

In the third stage, frame-wise detections are assembled into trajectories by solving a minimum-cost flow [50, 51] on a graph whose edges are weighted by spatial proximity and shape similarity and refined by “untangling” merges and splits [52]. This globally optimal assignment yields identity-consistent tracks through rapid motion and shape transitions. Complete trajectories are then analyzed to leverage shape evolution (via SI) for classifying event outcomes (e.g., productive vs. failed cargo transport) and for computing process-specific statistics (Fig. 1B). This integration of morphology with trajectory data underpins all subsequent analyses and provides a unified framework for outcome classification across diverse trafficking pathways at the PM. We validated the productivity labels against both a manually annotated dataset and orthogonal molecular markers of successful cargo delivery.

### Developing Shape2Fate for clathrin mediated endocytosis

CME is an ideal membrane trafficking process to showcase Shape2Fate and our shape descriptor, the Shape Index (SI). Live-cell TIRF-SIM used in this study provides ∼100 nm lateral resolution with an evanescent field depth of ∼100 nm (Supplementary Fig. 2). At this resolution, CCPs in RPE-1 cells with maximum diameters of ∼120-140 nm (peak-to-peak distance of the TIRF-SIM annulus; corresponding to ∼3-4 pixels) can be resolved as they progress through a characteristic sequence of morphologies [13]. We track CCPs across their lifetimes using fluorescently labeled clathrin light chain A (CLCa) [12, 13]. Early nascent CCPs appear as small, near-diffraction-limited disks corresponding to flat clathrin assemblies [53] present entirely in the TIRF plane (Fig 2A). As the clathrin coat assembles and the pit invaginates into a spherical dome, the TIRF-SIM signal typically becomes annular. This occurs because the curved periphery of the pit remains within the evanescent field while the central apex moves out of it, projecting the dome’s side-wall fluorophores as an annulus [12, 13]. Finally, as the budding CCP neck constricts, the TIRF-SIM signal reverts again to disk shape just before membrane scission of a fully formed vesicle [12].

**Figure 2.**
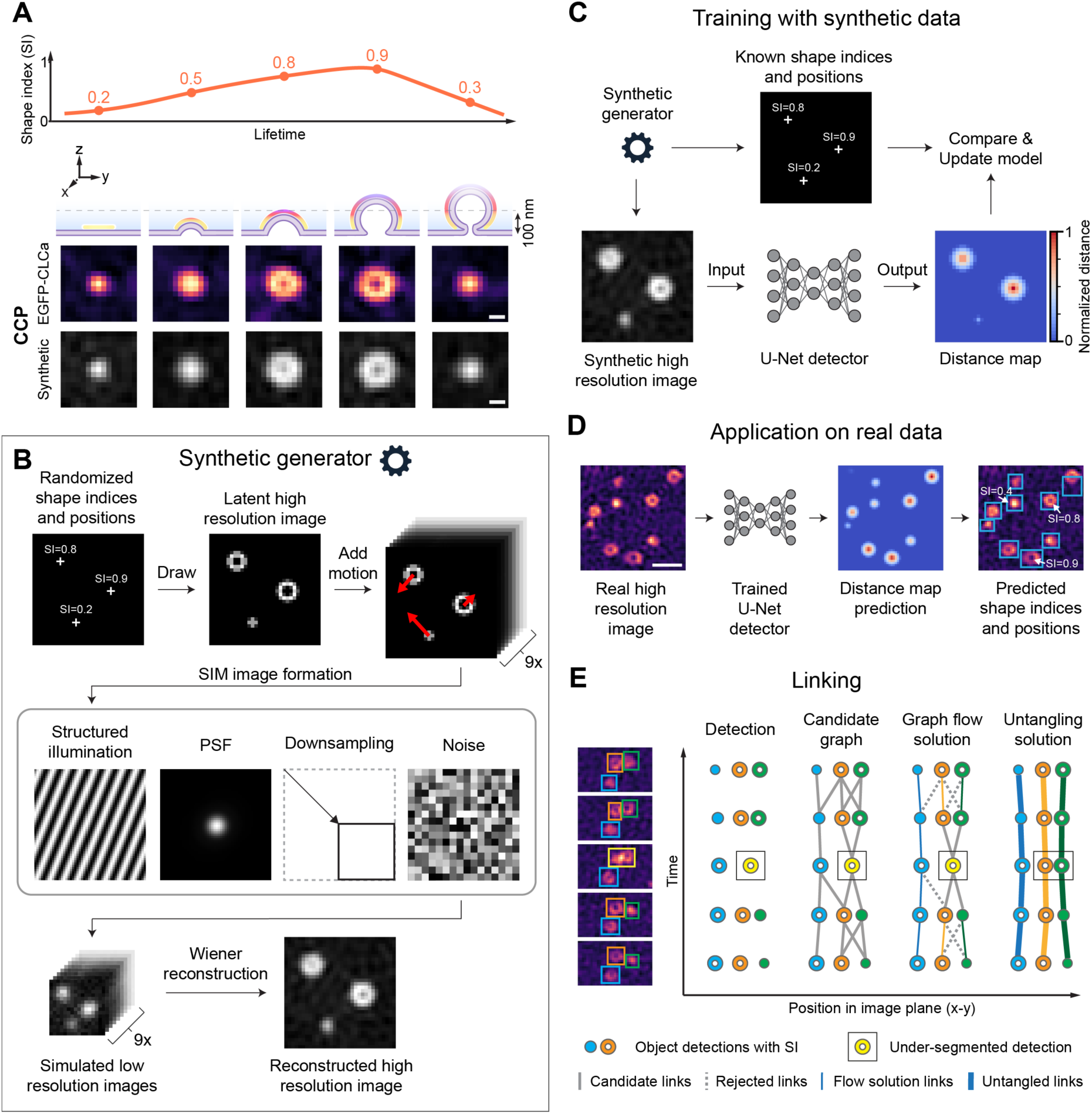
Shape Index-guided detection and linking of CCPs. **(A)** SI encodes CCP curvature progression over its lifetime (0 → disk, 1 → annulus). Top: SI trajectory with schematic morphologies. Bottom: representative real and synthetic CCP images at matched SI values. Scale bar: 100 nm. **(B)** Physics-based synthetic generator. Random SI values and positions are sampled to generate high-resolution CCP images; motion is added; SIM image formation is simulated (structured illumination, PSF, noise, downsampling) to produce nine raw frames, which are then Wiener-reconstructed into a high-resolution SIM image. **(C)** Training with synthetic data. Reconstructed synthetic images are fed to a U-Net that is trained to predict a normalized interior-distance map. Ground-truth positions and SI from the generator are used for supervised learning. **(D)** Application to real data. Reconstructed TIRF-SIM images are processed by the trained U-Net to provide distance maps; local maxima define object centres and their amplitudes encode per-object SI, yielding simultaneous detection and shape classification. Scale bar: 500 nm. **(E)** Linking and untangling. The Candidate graph shows all frame-to-frame associations in gray solid lines (edge cost = distance + |ΔSI|²). The Graph-flow solution retains optimal links (colored) and rejects others (gray dashed), forming provisional tracks. Untangling then removes biologically implausible merge/split patterns, yielding identity-consistent trajectories. The yellow node illustrates an under-segmented detection resolved by this process; over-segmentation and missed detections are handled analogously.

To quantitatively capture these morphological transitions, we implemented SI for CME as a continuous, normalized descriptor ranging from 0 to 1. Formally, the SI is here defined as the ratio between the apparent radius of the real CCP and the maximum possible CCP radius within the same dataset and cell line. Although expressed as a radius ratio, the SI implicitly encodes both shape and topological change: small radii correspond to diffraction limited disk-like structures, whereas larger radii reflect annular morphologies of the projected CCP dome. This definition effectively captures the disk-annulus transitions characteristic of a maturing CCP. Over its lifetime, the SI starts near 0, rises toward 1 as the pit invaginates, and drops at vesicle scission (Fig. 2A). Once the SI exceeds the threshold range, the pit has acquired sufficient curvature that progression to vesicle formation is strongly favoured and disassembly becomes rare. This interpretation is consistent with three independent lines of evidence. 3D single-molecule localization microscopy revealed a flat-to-curved transition at finite coat area [54]. Correlative electron and light microscopy showed that the onset of membrane bending marks a late, rarely aborted stage of CCP maturation [53, 55]. Thirdly, theoretical modelling predicts that vesicle formation proceeds cooperatively once a minimal local bending has been achieved [56, 57]. We therefore propose that SI provides a continuous and objective measure from a single clathrin channel for CCP progression, without requiring an additional curvature-specific or outcome-specific fluorescent reporter. This premise underpins the morphology-aware detection and productivity analysis enabled by Shape2Fate.

Building on this definition of SI for CME, we generated an extensive set of synthetic TIRF-SIM images of CCPs (∼ 0.5 million) spanning the full range of SI values to train our detection network (Fig. 2B, Supplementary Fig. 1A). Each synthetic image contains multiple CCP-like objects with known positions and assigned SIs. We represented each CCP in the latent high-resolution image by a 2D annular profile - the TIRF-SIM projection of a 3D dome - whose radius corresponds to its SI. We then simulated the TIRF-SIM imaging process (including structured illumination, optics, noise, and object motion; Fig. 2B; see Methods) to produce realistic high-resolution images with varied CCP appearances, arrangements and clustering (Supplementary Fig. 1A), all with ground-truth SI labels for supervised learning.

Using these annotated synthetic images, we trained a U-Net convolutional neural network [58] to detect CCPs and infer their morphology simultaneously (Fig. 2C). The network was trained to output a normalized interior-distance map for each frame, where every pixel value encodes its normalized distance to the nearest CCP boundary. In this representation, each CCP produces a local peak at its center with a value equal to its SI, tapering to zero at the edges and beyond. This distance-based encoding enables a single U-Net output that resolves adjacent CCPs and captures their underlying 3D morphological progression from the observed 2D TIRF-SIM projections. After training, we applied the U-Net to real TIRF-SIM time-lapse images (Fig. 2D). The predicted distance maps highlight candidate CCPs, with local maxima marking CCP centers and their amplitudes directly encoding the inferred SIs. Extracting these maxima enables simultaneous detection and morphological characterization of all CCPs within the field of view, without manual annotation. In theory, it would have been possible to train the network to perform detection directly on the nine raw frames, thereby effectively bypassing the SIM reconstruction step. However, to achieve sufficient robustness, it would need to handle all possible illumination-pattern configurations, resulting in a substantially more complex training procedure.

Next, detected CCPs in successive frames are linked into trajectories representing individual endocytic events (Fig. 2E). We formulate tracking as a graph optimization problem in which candidate links between detections over consecutive frames are scored based on both their spatial proximity and SI similarity (i.e., morphological continuity). This shape-aware linking cost favors consistent shape evolution along a track and helps prevent linking errors in crowded scenes. We also include auxiliary graph nodes that allow tracks to initiate or terminate and to handle occasional missed detections or spurious false positives by permitting track splits, merges, or skips. Solving a global minimum-cost flow problem [50] on this association graph yields a set of intermediate CCP tracks still containing biologically implausible splitting and merging detections. To eliminate these, we apply an “untangling” procedure [52] as a last refinement step. This step is itself an optimization problem that selects a consistent set of graph operations to enforce that each detection has at most one predecessor and one successor. The result is a collection of high-confidence non-conflicting CCP trajectories from initiation to completion or premature disassembly, each with a time series of SI values describing its shape evolution.

Shape2Fate leverages the shape evolution of each trajectory over time to classify the outcome of the CCP. Biologically, a productive endocytic event is expected to progress from a lower SI (nascent flat clathrin assembly) to a high SI (fully curved CCP) and then a rapid return to low SI values at scission. Abortive events, in contrast, exhibit incomplete or irregular SI trajectories, for example stalling at a low SI or dissipating before reaching a fully curved state (Supplementary Fig. 3). Operationally, for automated analysis we classify a trajectory as productive when it exceeds a determined SI threshold (described below), because reaching this high-curvature state most strongly predicts successful completion.

### Shape2Fate reliably identifies productive CCPs

We next asked whether the SI can reliably separate productive from abortive CCPs at the single-event level. To obtain an orthogonal molecular readout of productive CCP formation, we performed dual-color TIRF-SIM of SNAP-CLCa together with dynamin-2-mRuby3. Dynamin-2 is a large GTPase that assembles at late, high-curvature necks to catalyze membrane scission and exhibits a brief recruitment burst at productive events [59–61]. The dynamin burst therefore provides an orthogonal productivity marker, independent of clathrin intensity, for validating our SI-based CCP classification. To make the operation rule explicit and reproducible, we classified a CCP as productive if its SI exceeds 0.7 at any time point (Fig. 3Ai, Fig. 3B dotted line). This threshold was chosen to minimize classification error on a held-out, human-annotated validation set and corresponds morphologically to a high-curvature annular pit in TIRF-SIM (Fig. 3Ai), consistent with published evidence that curvature marks a commitment point for CCP maturation and scission [53–55].

**Figure 3.**
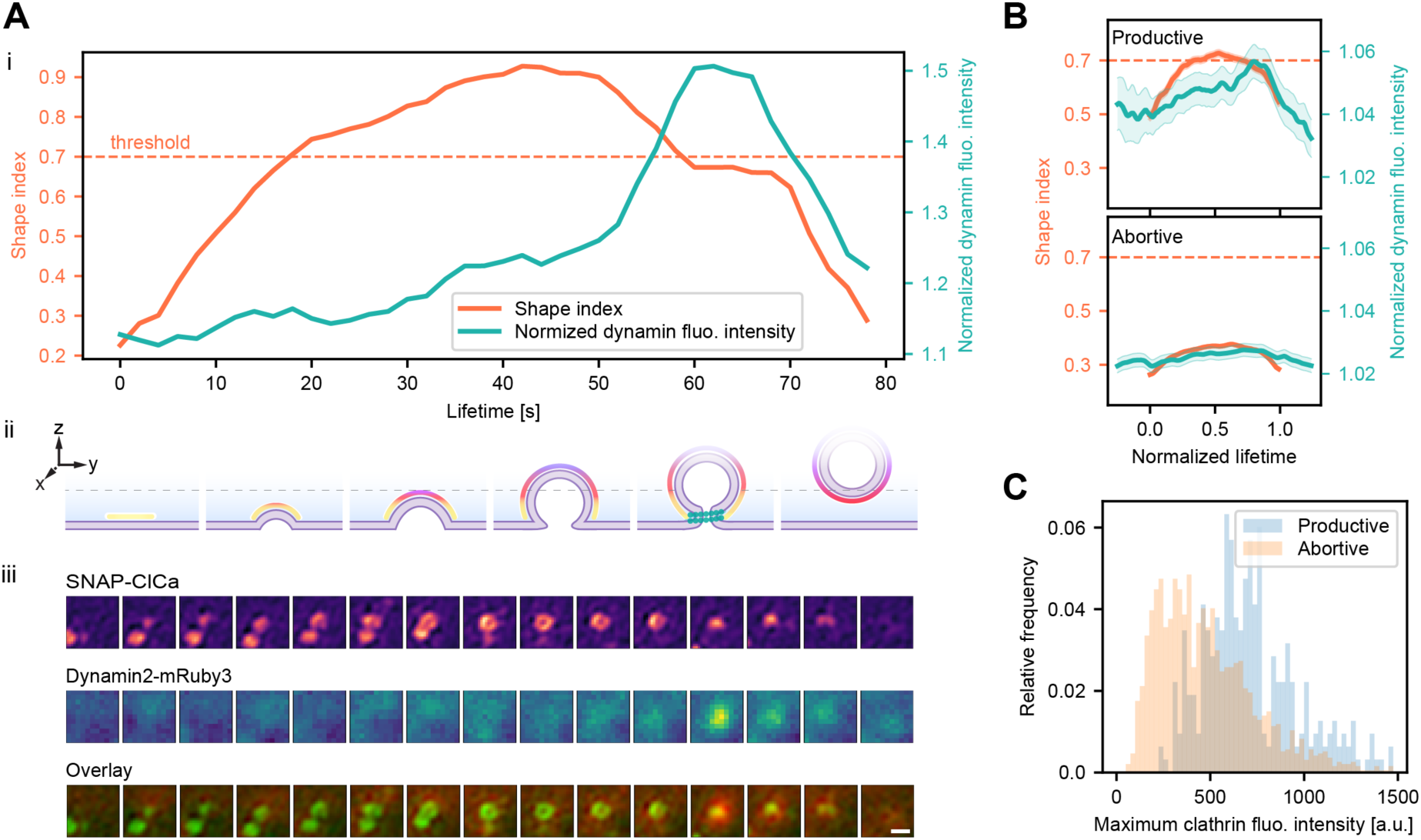
Orthogonal validation by dynamin-2 confirms that the Shape Index reliably identifies productive CCPs independently of fluorescence intensity. **(A)** Single-event example. (i) SI trace (orange) with productivity threshold at SI > 0.7 (dotted orange line) and dynamin-2 intensity (teal). (ii) Schematic contextualizing the SI metric: a side view of CCP maturation aligned to the SI function, with the dynamin-2 burst (teal) marking scission in SI-classified productive events. (iii) SNAP-CLCa and dynamin-2-mRuby3 snapshots across CCP lifetime. Scale bar: 250 nm. **(B)** Population validation. Time-normalized means (± SEM) of SI (orange) and dynamin-2 intensity (teal) for SI-classified productive (top, n = 478) and abortive assemblies (bottom, n = 2761). Productive events show a late dynamin burst coincident with SI decline, whereas abortive events lack this signature. Dotted orange line, productivity threshold at SI > 0.7 **(C)** Intensity versus SI. Distributions of maximum SNAP-CLCa intensity overlap substantially between SI-classified productive and abortive tracks, indicating that peak fluorescence does not reliably distinguish outcomes. All outcome labels in **B** and **C** derive from the SI criterion as shown in **Ai**.

Under this criterion, SI-classified productive events displayed a sharp dynamin-2–mRuby3 burst near the end of the lifetime that was temporally aligned with the SI drop expected at scission (Fig. 3A,B). In contrast, abortive SI-classified CCPs were largely dynamin-negative, indicating they do not progress to vesicle formation (Fig. 3B). These observations confirm that high-SI trajectories correspond to productive CCPs, while low-SI trajectories identify abortive assemblies.

This experiment also allowed us to compare our SI criterion to an intensity-only measure for productivity. As expected, the maximum SNAP-CLCa intensity showed substantial overlap between productive and abortive tracks (Fig. 3C), indicating that peak brightness alone is a weak discriminator of outcome. In contrast, SI provides single-event separation grounded in morphology rather than fluorescence intensity. Because SI is derived from shape, it is less sensitive than intensity thresholds to imaging conditions and expression levels. We demonstrate its generalisability by applying Shape2Fate to diverse cell types and acquisition settings across three microscope platforms (a custom-built eTIRF-SIM, Nikon N-SIM S, and DeltaVision OMX SR; see Methods).

### Shape2Fate achieves expert-level tracking

To benchmark Shape2Fate rigorously, we used three complementary evaluations. First, we performed a quantitative comparison against established automated CME tracking pipelines. Second, we evaluated performance relative to expert annotations by comparing Shape2Fate results with the inter-annotator range (human expert tracking variability). Third, we carried out a blinded paired comparison on tracks that experts had disputed. Because no tools currently exist for tracking subcellular membrane trafficking carriers at super-resolution, we chose cmeAnalysis [18], and TraCKer [12, 19] as the most functionally relevant and widely used TIRF-based baselines. Though developed over a decade ago, both remain the gold-standard tools for CCP tracking in conventional TIRF microscopy [12, 62]. These tools were designed to track diffraction-limited CCPs in conventional TIRF movies and they therefore fail to detect the complex morphologies resolved by TIRF-SIM. For this comparison, we applied the TIRF trackers to matched diffraction-limited inputs generated by averaging the nine raw pre-reconstruction frames, whereas Shape2Fate was evaluated on the corresponding reconstructed TIRF-SIM data (see Methods). Therefore, the performance differences demonstrated in Fig. 4 reflect two inseparable contributions: the two-fold resolution gain of TIRF-SIM and Shape2Fate’s shape-aware tracking, which exploit morphological information inaccessible to diffraction-limited methods.

**Figure 4.**
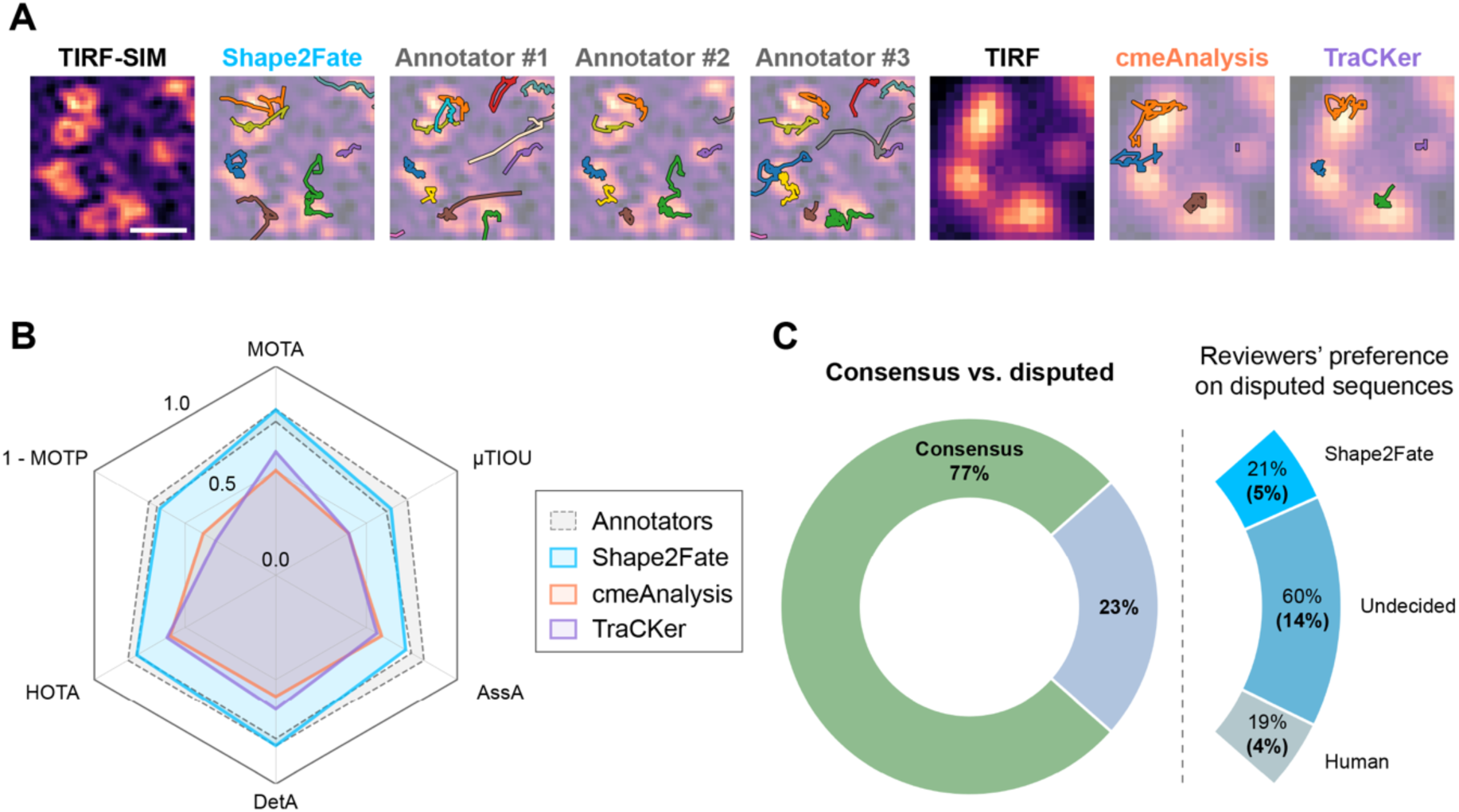
Benchmarking Shape2Fate against human annotators and TIRF trackers. **(A)** Qualitative comparison. Example sequence showing the TIRF-SIM image (left), Shape2Fate tracks and three human annotations (middle), and the matched TIRF frame with cmeAnalysis and TraCKer results (right). Trajectories are color-matched across panels. Scale bar: 500 nm. **(B)** Quantitative evaluation. Radar plot of MOTA, 1–MOTP, HOTA (DetA, AssA), and μTIOU (0–1; higher is better). The grey dashed band indicates the inter-annotator range (n = 3). Automated methods were parameter-tuned on the validation dataset before evaluation. **(C)** Blind preference on disputed sequences. Of all sequences, 77% reached human consensus and were not retested; the remaining 23% with disagreement among annotators were presented in a blinded, paired comparison (human vs Shape2Fate) to independent experts. On these disputed cases, reviewers preferred Shape2Fate in 21%, preferred the human annotation in 19%, and no unanimous preference was reached in 60%.

To benchmark this accuracy, we assembled a TIRF-SIM validation dataset annotated independently by three human experts. The dataset comprised ∼450 CCP trajectories from a typical RPE-1 cell, encompassing ∼13,000 individual detections per annotator. Because even expert human annotators disagree in their decisions when evaluating trajectories in dense clusters, and human tracking of individual CCPs in these circumstances becomes nearly impossible, this task lacks an absolute ground truth (Fig. 4A, Supplementary Movie 1). This ground truth challenge is not unique to this data domain [21]. To mitigate the lack of a single ground truth, we ran two complementary assessments. In the first, we assumed that individual expert annotations are noisy but that their inter-annotator range approximates the ground truth. In the second, we avoided assuming any single ground truth and instead used a blinded paired preference test to determine which method produces the most preferred results on the disputed, ambiguous cases.

For the first evaluation, we quantified tracking performance using six complementary metrics (Fig. 4B; see Methods). These metrics capture different aspects of tracking quality. Multiple object tracking accuracy (MOTA) [63] provides an overall reliability score by penalizing missed detections, false positives, and identity switches. Multiple object tracking precision (MOTP) [63] measures the localization error of matched detections; we report (1 − MOTP) so that higher values indicate better localization. Higher-order tracking accuracy (HOTA) [64] combines detection accuracy (DetA), which reflects how well true objects are identified, and association accuracy (AssA), which evaluates the correctness of temporal linking. Finally, we introduce here mean temporal intersection over union (μTIOU), a metric that quantifies the temporal continuity and overlap between predicted and ground-truth trajectories.

We first compared the three human annotators with one another to define the human inter-annotator range (Fig. 4B, gray dashed band), which serves as a practical reference rather than an absolute upper bound. Each automated method was then compared with each annotator and scored against this reference [21]. To ensure fairness, all methods (including Shape2Fate) were tuned on this dataset using their recommended procedures (see Methods), and we report their best performance achieved under that protocol. Shape2Fate scored ∼0.20 higher (i.e., a 20% absolute increase on the 0–1 metric scale) than the TIRF-based trackers, achieving performance within the range of variability observed between human annotators (Fig. 4B; Supplementary Table 1). An ablation study showed that including SI in the edge cost yields a ∼1–2% improvement in association metrics (HOTA, AssA, μTIOU). In TIRF-SIM, Shape2Fate separates and links individual CCP trajectories even within dense clusters that appear as merged spots in diffraction-limited TIRF - conditions that defeat manual tracking and intensity-based algorithms (Fig. 4A). On the validation dataset, we estimate that approximately 5% of CCP signal intensity cannot be adequately explained by the inferred CCP trajectories, corresponding to rare, highly complex or artifact-dominated clusters that remain challenging even for Shape2Fate (Supplementary Fig. 4 and Methods). This quantification demonstrates the method’s ability to handle most difficult cases. Strong performance also generalized to a separate CME testing dataset acquired on a different TIRF-SIM imaging platform and under different acquisition settings (438 CCP trajectories; ∼35,000 detections; Supplementary Table 2).

Finally, we directly compared Shape2Fate with human annotators on the most challenging and ambiguous sequences. We conducted a blinded paired preference test focusing on the ∼23% of movie sequences in which the three annotators had originally disagreed, that is, cases with no clear reference answer even among experts (Fig. 4C, left). Independent expert reviewers evaluated complex CCP tracks in clusters, comparing two anonymized track sets (one human, one Shape2Fate, identity hidden) and choosing which better preserved object identity and motion continuity. This head-to-head test confirmed that Shape2Fate achieves human-level performance on difficult CME cases. Reviewers slightly favored Shape2Fate tracks in 21% of cases over the human annotator’s (19%), and in most comparisons (60%) they had no unanimous preference. Moreover, outside of these complex cases Shape2Fate had already matched the consensus of human annotators in the remaining 77% of sequences, meaning it agreed with annotator calls whenever the annotators themselves concurred. Taken together, this three-tiered validation (i.e. benchmarking against established CME trackers, calibration against human experts, and blinded preference testing on disputed sequences) demonstrates that Shape2Fate reaches human-expert accuracy across a broad spectrum of tracking difficulty.

### Shape2Fate extends to exocytosis, resolving pleomorphic carriers and their fusion events

Having established and validated Shape2Fate for CME, we next asked whether the same shape-based logic could tackle a more challenging regime of exocytosis. Exocytic carriers are highly pleomorphic, ranging from elongated tubules to spherical vesicles, and exhibit markedly different arrival, docking, and fusion kinetics. To our knowledge, pleomorphic tubular exocytic carriers have not previously been tracked quantitatively at the single-carrier level, in part because conventional TIRF-based tracking algorithms are primarily tailored to near-spherical, diffraction-limited spots [17, 20, 65]. To address this challenge with Shape2Fate, we retained the overall pipeline structure (including SIM reconstruction, detection, linking, and quantitative analysis) and introduced task-specific changes for exocytosis.

We applied Shape2Fate to two exocytic systems that differ fundamentally in regulation, carrier morphology, and fusion signature: a synchronized constitutive secretion assay producing pleomorphic tubular carriers with visible cargo throughout trafficking, and insulin-regulated GSVs that are spherical and detectable only upon pH-dependent fusion. First, we used engineered SH-SY5Y cells with RUSH, a timed-release model in which a small-molecule cue (biotin) synchronizes delivery of carriers with a transmembrane reporter to the PM [37]. Our transmembrane reporter (mEmerald–LAMP1) has an extracellular fluorescent tag and was engineered to avoid rapid re-internalization, enabling a clean readout of arrival and fusion [38]. After biotin stimulation, cargo reached the surface in pleomorphic tubular carriers: many fused with the PM, whereas others remained highly mobile without docking or fusion. Second, we tested Shape2Fate on physiologically relevant insulin-stimulated GLUT4 exocytosis in adipocytes, where insulin signalling promotes the translocation of GSVs to the PM [66]. Together, these models challenge Shape2Fate across distinct carrier shapes (tubules versus approximately spherical vesicles), fluorescence intensity profiles, and timescales, testing whether a single shape-aware pipeline can detect, link, and classify exocytic events across systems. In TIRF-SIM 2D projections, RUSH carriers appear either as elongated tubules when moving in-plane or as small compact disks (2D projections of tubules perpendicular to the membrane). Successful fusion yields an annular fusion pore followed by a transient cargo burst and lateral diffusion (Fig. 5A, top). In contrast, vesicles containing luminally tagged pHluorin–GLUT4 are largely dim before fusion. Fusion produces a sudden, high-contrast pHluorin fluorescence flash at the PM (Fig. 5A, bottom) [67].

**Figure 5.**
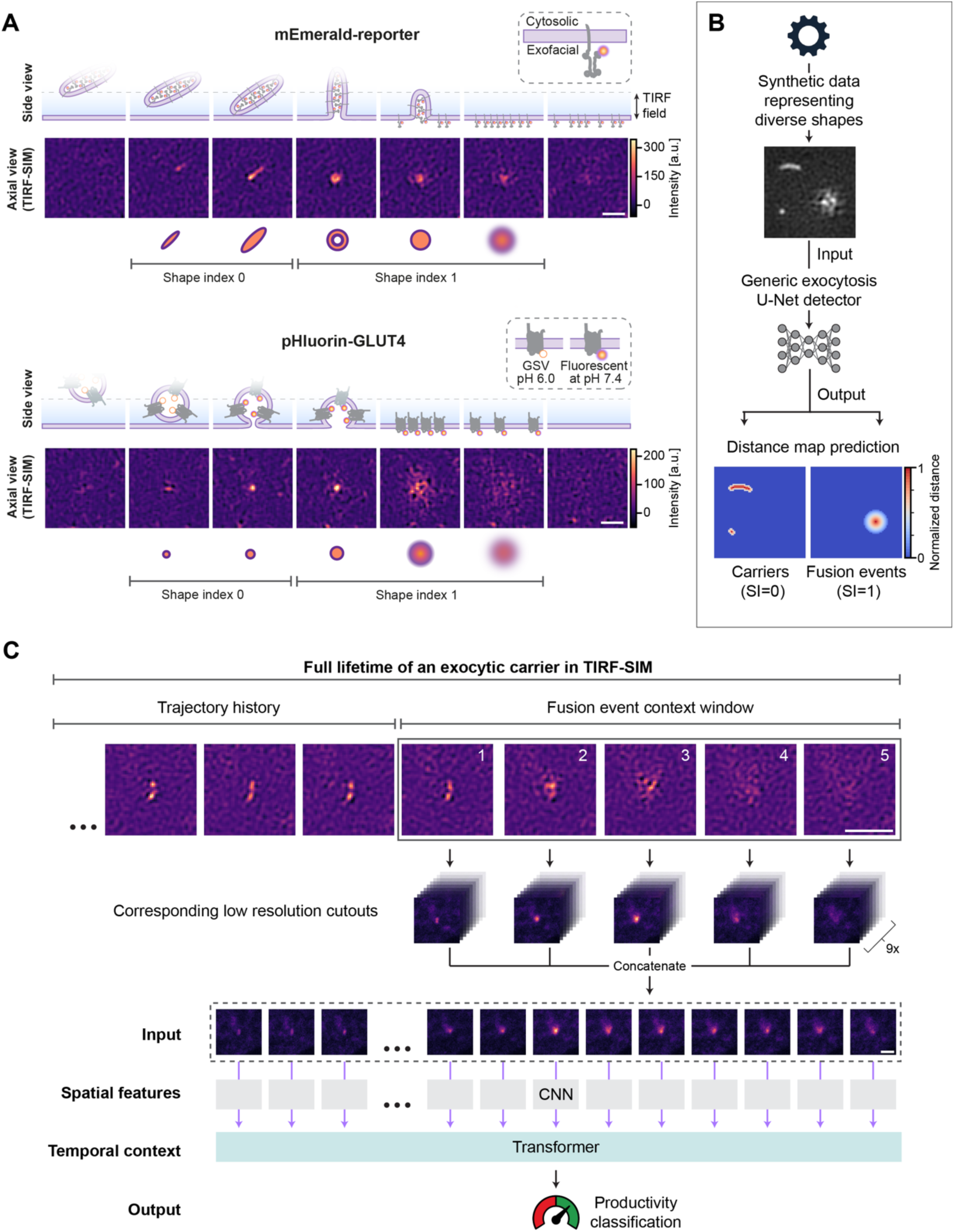
Extending Shape2Fate to morphologically diverse exocytic carriers. **(A)** Cargo-labeled exocytic events in two distinct systems. Side-view schematics and axial TIRF-SIM frames for mEmerald-reporter (RUSH) and pHluorin-GLUT4 illustrate arrival, docking, pore opening, and cargo release. Approaching carriers appear as disks or elongated tubules and are assigned SI = 0; pore opening produces a transient annulus with subsequent diffusion, assigned SI = 1. Inset-top: schematic of the mEmerald-reporter construct showing cytosolic and exofacial domains. Inset-bottom: pHluorin-GLUT4 and pH-dependent fluorescence. Scale bar: 500 nm. **(B)** Core detector trained on synthetic data. A physics-based generator produces diverse carrier shapes and fusion flashes. A U-Net is trained to predict two normalized interior-distance maps: carriers (SI = 0) and fusion events (SI = 1). Connected-components analysis of these maps provides simultaneous localization and class labels. **(C)** Auxiliary fusion classifier. For each candidate fusion, the raw TIRF stacks corresponding to the final five TIRF-SIM frames are concatenated and analyzed frame-by-frame by a shared CNN to extract spatial features, which are then fed as a sequence into a transformer that captures temporal context. The classifier outputs a binary fusion/non-fusion decision, suppressing artifact-driven false positives. Scale bar: 1 µm.

Because exocytic carriers exhibit substantially more diverse projected geometries than the ordered curvature-dependent progression observed in CME, we retrained the shape-aware detector within the Shape2Fate framework (Fig. 5B) using a new synthetic dataset (Supplementary Fig. 1B). These simulations span spherical vesicles, tubular carriers, and fusion events represented as sudden isotropic signal bursts that mimic cargo release and lateral diffusion. This adaptation enabled Shape2Fate’s consistent detection, linking, and classification of exocytic events across both systems within a unified, SI-guided framework.

Building on the same detection-linking core of the pipeline, we revised the shape classification scheme to accommodate the abrupt signal changes at fusion. Instead of a continuous SI (0–1) used for CCPs, exocytic detections were assigned a binary SI: 0 for intact vesicles or tubules, and 1 for productive fusion events. The U-Net detection network was accordingly modified to output two separate distance maps - one for each class - thereby simultaneously localizing carriers and flagging fusion occurrences (Fig. 5B). Because exocytic carriers span more irregular shapes and transitions than CCPs, we replaced the peak-based detection in the distance maps used for CME with connected-component segmentation of the distance maps. The centroid of each segmented object defined its location, and its class (0 or 1) was determined by which map it originated from. Detections were linked with the same graph-based algorithm, accounting for faster motion and sudden shape changes, e.g., carrier tracks ending in a single-frame fusion flash (Supplementary Fig. 5).

Because exocytic fusion and the initial high intensity cargo burst occur on a sub-second timescale, fusion often appears as a single-frame flash that can induce motion-like SIM reconstruction artifacts. These artifacts occasionally cause the detector to misclassify spurious intensity spikes as fusions. To suppress such false positives, we incorporated an auxiliary fusion classifier that examines the raw unreconstructed low-resolution images corresponding to the last five TIRF-SIM frames for each candidate event, independently of prior tracking (Fig. 5C). The model couples a shallow CNN [68] for spatial features with a transformer [69] for temporal dynamics and outputs a binary fusion/non-fusion decision based on the carrier’s spatiotemporal signature. Unlike the core detector, which operates on individual reconstructed high-resolution frames and can therefore be trained entirely on synthetic snapshots, the auxiliary fusion classifier must evaluate a temporal sequence of low-resolution frames capturing carrier approach, docking, and fusion. Faithfully simulating such sequences would require pathway-specific biological priors, such as carrier motility regimes, dwell-time distributions, and fusion-pore kinetics. Our synthetic generator intentionally avoids these assumptions and instead reproduces only the carrier appearance in single reconstructed TIRF-SIM snapshots. To circumvent this, the fusion classifier was trained on real annotated fusion events with mEmerald-reporter in RUSH engineered SH-SY5Y cells or pHluorin-GLUT4 in insulin-stimulated adipocytes to learn features common across exocytic systems, cell types, and imaging conditions. We intentionally favor sensitivity during detection and linking, then use the classifier as a secondary filter to retain only high-confidence fusions and suppress artifact-driven false positives.

Next, we evaluated Shape2Fate exocytosis tracking on an expert-annotated RUSH test set. Shape2Fate tracked the arrival and fusion of most annotated exocytic carriers (745 carrier trajectories; ∼25,000 detections; Supplementary Table 3, Supplementary Movie 2). To our knowledge, Shape2Fate is among the first approaches to quantitatively follow pleomorphic exocytic carriers near the PM over extended lifetimes, that include both carrier arrival and fusion. Benchmarking tracking metrics for exocytosis were lower than those for CME (Supplementary Tables 2–3). In the exocytic regime we deliberately favor tracking sensitivity, so Shape2Fate also captures low-intensity and very short-lived fusion traces that human annotators typically omit. When benchmarked against these human annotations, such additional Shape2Fate tracks were conservatively scored as false positives. Although this design choice slightly lowers apparent precision, it ensures more comprehensive sampling of exocytic events for downstream quantitative analyses. For comparison, most existing approaches focus only on detecting the diffraction-limited terminal fusion flash rather than comprehensively tracking arriving carriers throughout their lifetime to fusion [65, 70, 71]. Because IVEA [72] is the most recent method for detecting exocytic fusion events, we used it as a benchmark for comparison with Shape2Fate. Of note, both IVEA and our fusion classifier operate on TIRF-resolution input. Under these matched conditions, our fusion classifier substantially outperformed the pre-trained IVEA model for productive exocytosis classification (Supplementary Table 4).

In summary, the SI-guided framework transfers across the multiple membrane-trafficking pathways tested here, accommodating distinct carrier dynamics, morphologies, and fusion signatures to reconstruct trajectories in both endocytosis and exocytosis. This flexibility is especially useful for exocytosis, where Shape2Fate supports both pH-sensitive reporters and constitutively fluorescent markers because event classification is not tied to a single tagging strategy.

### Shape2Fate reveals exo–endocytosis coupling in synchronized constitutive secretion on global and local scales

We next asked whether endocytic and exocytic events are actively coordinated in space and time. Shape2Fate enabled us to address this directly, as it tracks both processes simultaneously at event-level resolution (∼100 nm) within the same cell. Using the biotin-triggered RUSH assay to synchronize mEmerald–reporter exocytosis in SH-SY5Y cells, we acquired a large dual-channel TIRF-SIM dataset capturing exocytosis alongside CME through interleaved imaging (Fig. 6). Shape2Fate reconstructed trajectories for ∼2,000 exocytic fusion events and ∼100,000 endocytic CCP events across 42 cells, providing an extensive collection to quantify both local and global coupling between these membrane-trafficking pathways.

**Figure 6.**
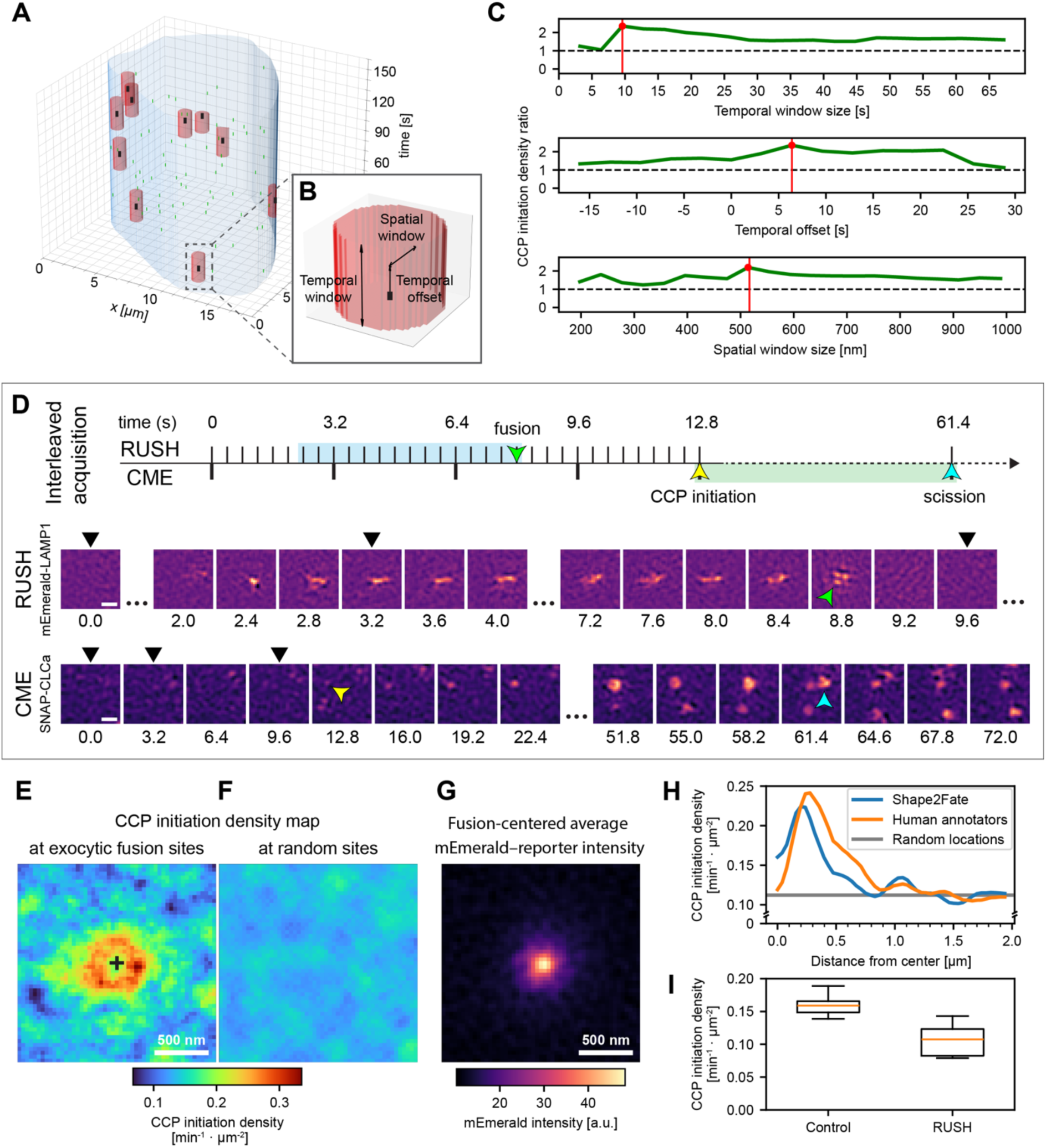
Shape2Fate reveals local and global scales of exo-endocytic coupling during constitutive secretion. **(A)** Schematic of the approach used to quantify the initiation of CCPs near productive exocytic events. The translucent blue volume (cell mask) outlines the region occupied by the cell throughout the recorded time-lapse. Green dots mark CCP initiation events, while red dots indicate productive exocytic fusion sites. Red cylinders highlight spatiotemporal regions around each fusion site, with their size and duration selected by an exploratory parameter sweep. **(B)** Magnified view of one spatiotemporal region with its adjustable parameters highlighted. **(C)** Line plots show the CCP-initiation density ratio (local/background) as each parameter is swept. Red markers indicate the settings used in our analysis: 0-15 s window starting at fusion and ∼500 nm radius. **(D)** Interleaved acquisition strategy used to capture the temporal relationship between exocytosis (RUSH mEmerald-reporter) and CME (SNAP-CLCa). Top: Acquisition timeline showing rapid imaging of the RUSH exocytosis channel (frames every 0.4 s) interleaved with CME imaging intervals (frames every 3.2 s). Arrowheads mark key events: fusion of a LAMP1-positive carrier (green), initiation (yellow), and scission of a CCP (cyan). Bottom: Representative TIRF-SIM image sequences from both channels illustrating coordinated events. Black arrowheads indicate corresponding frame times. Scale bar: 250 nm. **(E)** Heatmap of CCP-initiation density centered on exocytic fusion. Averaging within the 500 nm and 0-15 s window around productive fusion sites reveals a pronounced peri-fusion hotspot (n > 1,000). Scale bar: 500 nm, the white dotted line in the color bar represents 0.23 min⁻¹ µm⁻². **(F)** Averaging CCP-initiation density around random positions (n > 10,000) produces a uniform CCP initiation density heatmap without a hotspot. **(G)** Averaged mEmerald-reporter intensity centered on the sites of productive exocytic fusion events (n > 1,000), aligned to the time of fusion. Scale bar: 500 nm. **(H)** One-dimensional radial profile of the data from previous panels, plotting mean CCP initiation density as a function of distance from productive exocytic events. Shape2Fate detections (blue) closely match human annotations (orange) and the grey line shows mean CCP initiation density at random positions. **(I)** Comparison of global CCP initiation densities in control cells and cells under synchronized constitutive secretion (p = 0.0001, n = 16 cells).

To test for local coupling first, we defined a cylindrical spatiotemporal region around each productive fusion event and measured the rate of CCP initiation within that region relative to the rest of the cell (Fig. 6A, B). We swept spatial radius, temporal duration, and temporal offset of the cylindrical window relative to each exocytic event. Line plots from this exploratory sweep show how the mean ratio of local-to-global CCP initiation density varies with each parameter (Fig. 6C). This analysis identified a peak within a spatiotemporal window with ∼500 nm radius and 0-15 s after the fusion, where CCP-initiation density was more than double the global baseline (Fig. 6C, D). To visualize this phenomenon in space, Fig. 6E displays a heatmap of CCP initiation occurring up to 15 s after and spatially around each productive fusion event in the RUSH system, with brighter colors representing higher CCP initiation density. This heatmap centered on exocytic fusion sites reveals a pronounced annular hotspot of CCP initiations. Using the mEmerald–reporter intensity maximum to define spatially the fusion (Fig. 6G), the CCP-initiation density reaches ∼0.23 min⁻¹ µm⁻² at ∼250 nm from that point, as seen in the corresponding one-dimensional radial profile of the heatmap (Fig. 6H). This distance is comparable to the sum of the effective radii of the exocytic patch (Fig. 6G) and a nascent CCP and indicates the spatial precision of our detection and linking (Fig. 6D-H). About one-quarter of productive fusion events were closely followed by initiation of one or more CCPs (Supplementary Fig. 6, Supplementary Movie 3). Manual human annotation provided an independent verification that CCP initiation was significantly enriched around recent fusion sites (Fig. 6H).

Although more CCPs initiated near fusion sites, their maturation efficiency, measured as the ratio of productive to abortive events, matched the cell-wide baseline. This suggests that the coupling mechanism primarily biases where CCPs form rather than how efficiently they mature once initiated (Supplementary Fig. 7). To test whether synchronized exocytosis alters endocytic activity at the global, cell-wide level, we quantified CCP initiation densities before and after biotin addition in the SH-SY5Y cells engineered with RUSH. The global CCP initiation density fell by ∼25% (from ∼0.16 to ∼0.11 min⁻¹ µm⁻²; Fig. 6I), which, if considered alone, would suggest that massive exocytic flux broadly suppresses CME. Thus, rather than simply suppressing endocytosis, the synchronized exocytic burst triggers a spatial redistribution of endocytic capacity: CCP initiation doubles within ∼500 nm of each fusion site, while declining ∼25% across the rest of the cell. Shape2Fate therefore resolves the apparent global downregulation into a targeted local reallocation of endocytic machinery to sites of membrane influx.

### Shape2Fate reveals that adipocytes employ an inverted coupling logic for GSV fusions

To investigate exocytic-endocytic coupling in a physiologically regulated system, we next applied Shape2Fate to insulin-stimulated 3T3-L1 adipocytes expressing pHluorin-GLUT4 and SNAP–CLCa. As a prerequisite, we confirmed successful differentiation and stimulation by monitoring bulk PM accumulation of GLUT4-pHluorin over 40 min using TIRF microscopy in day-nine differentiated adipocytes after insulin addition (Supplementary Movie 4). TIRF-SIM imaging revealed that adipocyte CCPs were on average ∼1.5× larger in diameter than those in RPE-1 cells (Fig. 7A), a difference confirmed by EM (Fig. 7B). To account for this larger pit size, we expanded the initial synthetic generator to cover CCP diameters up to 250 nm and adjusted the shape-index radius accordingly. This adaptation ensured that the detection model is robust across all CCP sizes encountered in all the tested cell lines. With these adaptations in place, Shape2Fate identified ∼3,000 GSV fusion events alongside ∼50,000 CCP trajectories in 23 cells by dual-colour TIRF-SIM time-lapse series (Supplementary Movie 5). Analysis of these trajectories revealed a coupling architecture distinct from that of the synchronized constitutive secretion in RUSH assay. Whereas synchronized constitutive secretion nucleates de novo CCPs, insulin-stimulated GSV fusions in adipocytes instead target pre-existing CCPs as preferred docking sites. This represents an inverse spatiotemporal hierarchy in which endocytic machinery spatially precedes exocytosis rather than following it (Supplementary Movie 5-6). A representative dual-channel time-lapse illustrates this coupling: a GLUT4 fusion event occurs at a membrane site already occupied by a CCP, and the released GLUT4 co-localises with the pit within seconds (Fig. 7C, Supplementary Fig. 8, Supplementary Movie 6). Shape2Fate quantifies this spatial enrichment at multiple scales. CCP density in the peri-fusion window is ∼86% higher than baseline (Supplementary Fig. 9), and ∼30% of fusion events occur within 500 nm of a pre-existing CCP (Supplementary Fig. 10). Overall, fusion-to-CCP distances are significantly shorter than expected by chance (Fig. 7D–F). This spatial interaction was highly non-random (p=2e-5, >0.916 with 95% conf., one-sided one-sample t-test, n=3, 16 cells), consistent with preferential targeting of GSVs to pre-formed endocytic sites. By delivering GLUT4 directly into CCP-rich hotspots, cells may partially bypass reliance on passive diffusion. Approximately 33% of the pHluorin–GLUT4 fluorescence intensity appearing at the surface upon vesicle fusion quickly became associated with a nearby (<500 nm) CCP within ∼2–3 seconds (Fig. 7G). Consistent with this, GLUT4 intensity at the PM was higher within CCPs than outside them (Supplementary Fig. 11). This post-fusion recruitment was most pronounced under insulin stimulation, suggesting that insulin not only triggers GLUT4 exocytosis but also promotes spatial coordination between GSV fusion sites and CCP locations (Fig. 7G, Supplementary Fig. 8). Together, these findings demonstrate that in insulin-responsive adipocytes, regulated exocytosis is coupled to the CME machinery, a relationship that Shape2Fate quantified to a previously inaccessible level of spatiotemporal resolution (Fig. 8).

**Figure 7.**
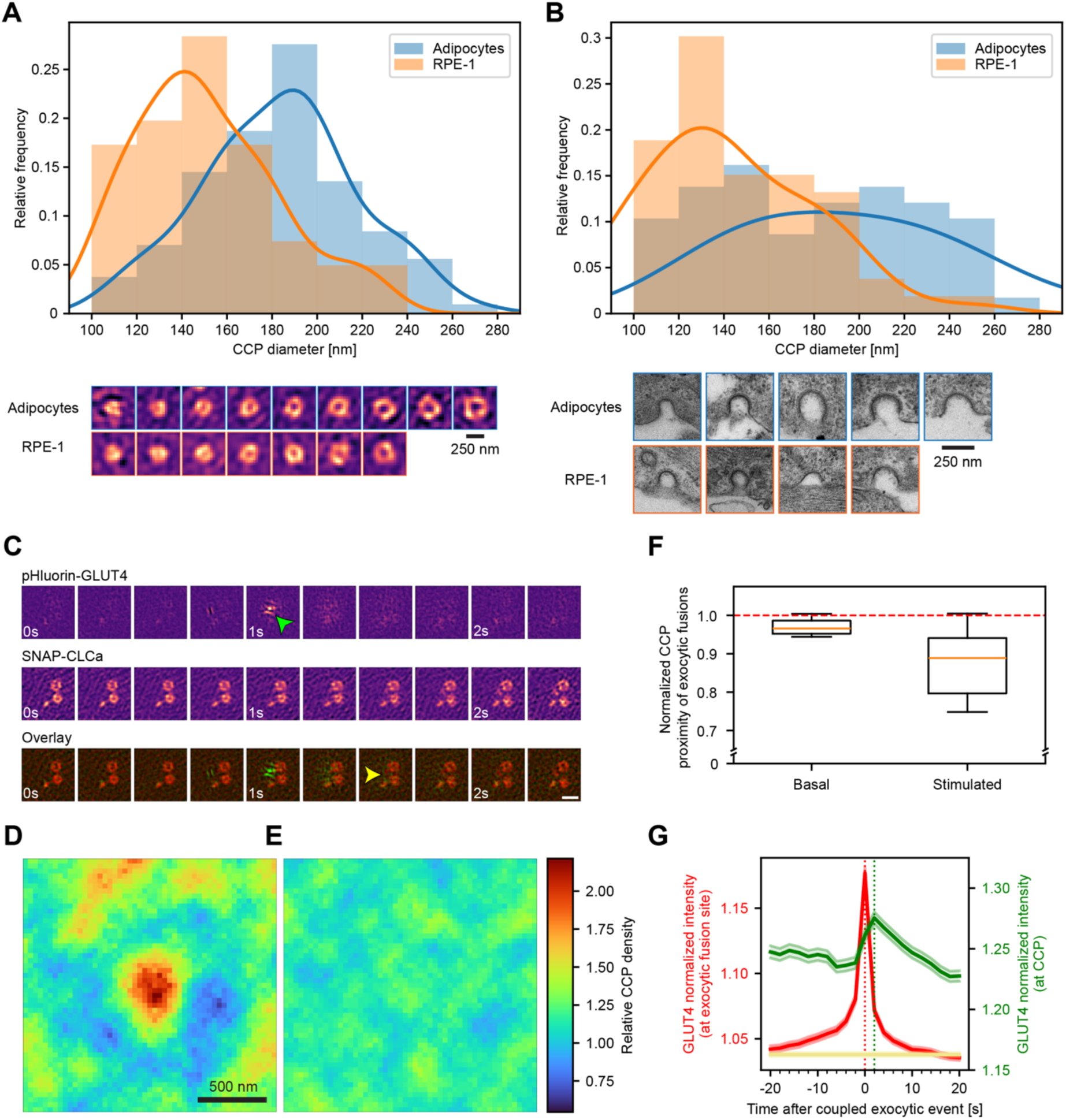
Shape2Fate reveals that pre-existing CCPs direct GSV fusion sites in adipocytes, enabling seconds-scale GLUT4 capture. **(A)** Distribution of CCP radii measured by Shape2Fate in adipocytes (blue) and RPE-1 cells (orange); below, representative TIRF-SIM cut-outs, n = 3, 20 cells. **(B)** Histogram of CCP diameters measured from EM images of adipocytes (N = 58, 5 cells) and RPE-1 cells (N=52, 7 cells). **(C)** Dual-channel time-lapse snapshots of a single GSV fusion aligned to pre-existing CCPs. Top: pHluorin-GLUT4 shows the fusion burst (green arrowhead). Middle: SNAP-CLCa reveals a CCP at the same membrane region. Bottom: overlay highlights rapid co-localization (yellow arrowhead) within seconds of fusion. Scale bar: 500 nm. **(D, E)** Heatmaps of CCP density. Scale bar: 500nm. Fusion-centred averaging reveals a pronounced CCP-enriched hotspot around productive exocytic sites **(D)** compared to random control positions **(E)**. **(F)** Comparison of the relative mean distance from GSV fusion sites to the nearest CCP under basal and insulin-stimulated conditions, plotted relative to a random expectation (red dashed line), p = 0.001 **(G)** Time-aligned averages centred on fusion (t = 0): mean pHluorin-GLUT4 intensity at the fusion site (red); concomitant pHluorin-GLUT4 at nearby CCPs (green), pHluorin-GLUT4 intensity at CCPs 1 µm distant from any GSV fusion site (yellow). Data presented from n=3 independent replicates, with at least 3 cells per condition.

**Figure 8.**
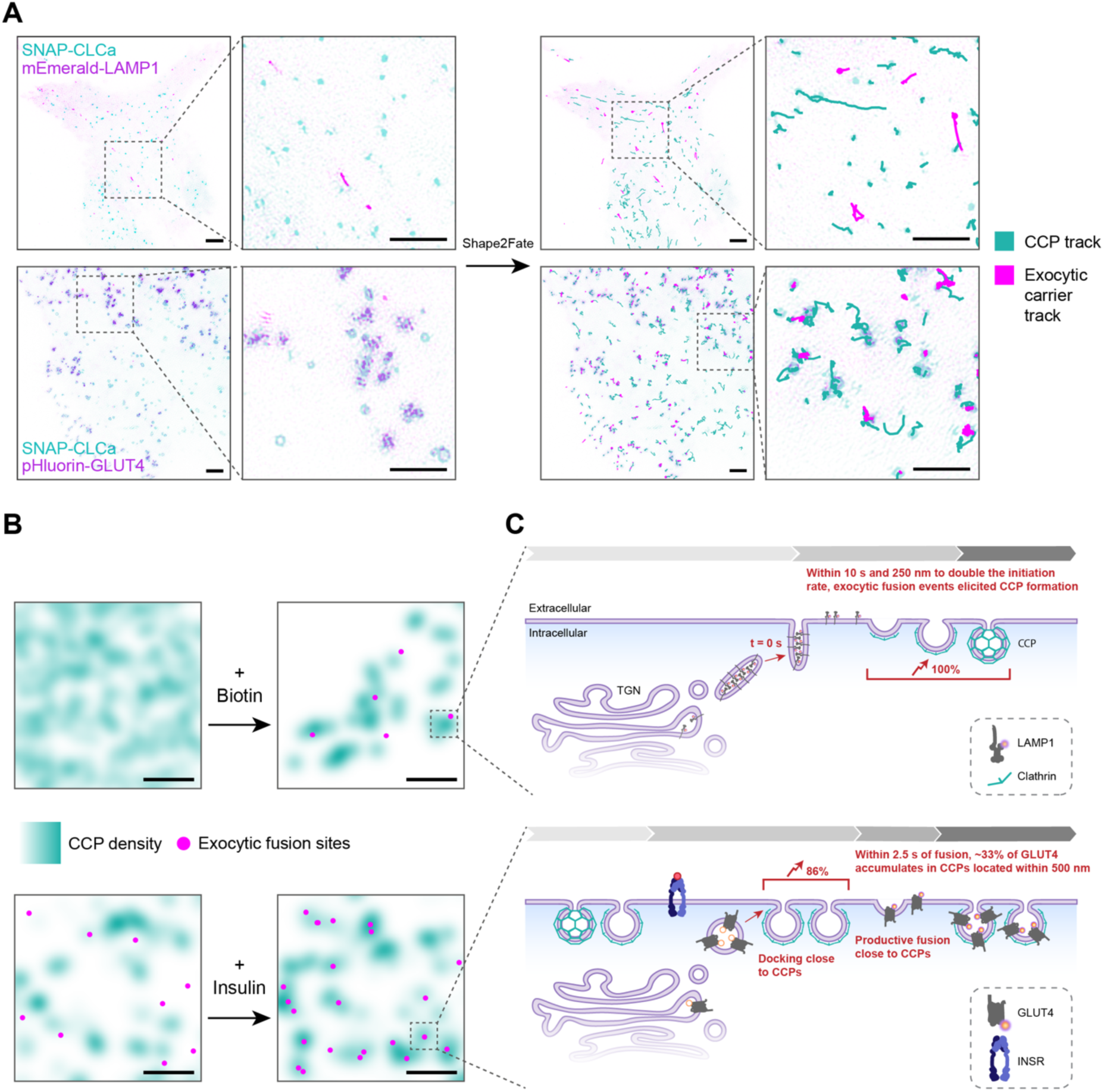
Shape2Fate maps two diametrically opposed architectures of exo-endocytic coupling. **(A)** Dual-channel TIRF-SIM images (left) and Shape2Fate tracking output (right) for synchronized constitutive secretion (top) and insulin-stimulated adipocytes (bottom), illustrating the framework’s ability to resolve both systems despite their contrasting carrier morphologies and CCP size distributions, densities, and spatial arrangements. Scale bars: 2 µm. **(B)** Spatial maps of CCP density (teal) and exocytic fusion sites (magenta dots) before and after stimulation with biotin (synchronized constitutive secretion with RUSH, top) or insulin (adipocytes, bottom). Scale bars: 2 µm. **(C)** Integrative model of the two coupling architectures resolved by Shape2Fate (quantitative details in Fig 6-7). Top: RUSH fusion nucleates de novo CCPs locally. Bottom: adipocyte GSVs fuse at pre-existing CCPs, enabling rapid cargo capture.

## Discussion

To our knowledge, Shape2Fate provides the first unified, automated pipeline for event-resolved tracking of both endocytic and exocytic carriers at super-resolution within the same cell. A fundamental motivation for Shape2Fate’s geometry-first design is that intensity-only criteria for detection, linking and classification are fragile in practice. They depend on arbitrary thresholding and are sensitive to acquisition-dependent fluctuations arising from low SNR, photobleaching and tracking artefacts. More sophisticated intensity-derived metrics can improve classification in some settings, for example the clathrin-coat assembly growth rate [18, 19] and the disassembly asymmetry score (DASC) [8]. However, approaches such as DASC require very large numbers of traces per condition (>10^5) to stabilize threshold-free classification, which is not always feasible experimentally. In contrast, by leveraging geometry as the primary signal, Shape2Fate yields robust performance across the various TIRF-SIM platforms, acquisition settings, and membrane-trafficking pathways tested here. Importantly, the same morphology-aware framework applies to both endocytic and exocytic carriers, because these membrane-trafficking carriers undergo characteristic shape transitions that reflect their progression toward distinct functional outcomes. More broadly, the geometry-first principle for tracking and outcome classification, introduced in this work, may extend beyond membrane trafficking to any cellular process in which structures pass through a reproducible morphological sequences, for example, autophagosome biogenesis or mitochondrial fission at ER contact sites.

The robustness of Shape2Fate’s geometric approach addresses a regime that is currently under-served by existing general-purpose MOT methods. Recent MOT frameworks such as Ultrack [22] and Trackastra [23] are powerful solutions for cells or nuclei that can be segmented reliably and are large enough to support overlap-based measures (e.g., intersection-over-union) or stable region descriptors [21]. Membrane-trafficking carriers imaged by TIRF-SIM, in contrast, are nanoscale, fast-moving, and often lack well-defined segmentation boundaries, especially when they rapidly change morphology over time, making segmentation-first pipelines unreliable [21]. Shape2Fate therefore uses an implicit representation that encodes both position and shape (via a distance-map–based encoding), and it performs global linking with graph optimization using link costs that integrate spatial proximity with differences in a shape descriptor (SI). This combination makes it possible to track sub-diffraction carriers robustly even when explicit object masks are ambiguous.

Applying Shape2Fate to two mechanistically distinct secretion systems exposes a set of organizing principles for how non-neuronal cells spatially coordinate membrane trafficking. The most striking biological finding is that the two systems, synchronized constitutive secretion and insulin-stimulated GLUT4 exocytosis in adipocytes, display diametrically opposed coupling architectures. In the synchronized constitutive secretion, exocytic fusion of mEmerald-reporter carriers is rapidly followed by de novo nucleation of CCPs near the fusion site (Fig. 8C, top). Because this reporter lacks obvious adaptor-binding motifs, the local recruitment of CME machinery is unlikely to be driven solely by direct cargo - adaptor interactions. Two plausible and potentially synergistic mechanisms may account for this local enrichment of CCP formation. First, exocytic fusion occurs preferentially at PI(4,5)P₂-enriched plasma membrane microdomains [73], and PI(4,5)P₂ is also actively generated on arriving carriers by Arf6-dependent PIP5K1C activity [74]. Second, fusion locally adds membrane area and reduces tension, lowering the energy barrier for CCP curvature generation. Concurrently, global CCP formation drops by ∼25%, revealing that de-novo CCP formation at fusion sites operates under a capacity constraint: increased endocytic activity at one site comes at the expense of another. A bulk assay such as transferrin uptake, or any quantitative approach that averages over the entire cell surface, would capture only this global decrease and conclude that exocytosis suppresses endocytosis. Event-level spatiotemporal analysis, as Shape2Fate provides, reveals instead a spatial reallocation of endocytic capacity toward sites of membrane influx.

This coupling hierarchy also differs from coupling mechanisms described in other secretory systems. In INS-1 cells, vesicle fusion has been reported to engage CME via a “hopping” mechanism in which pre-existing CCPs relocate to fusion sites [75], while in Drosophila salivary glands, new clathrin coats assemble directly on the large crumpled membranes of secretory vesicles [76]. The RUSH exo-endo coupling also differs fundamentally from the well-characterized coupling at neuronal synapses. At synapses, exo-endocytic coupling is enforced by permanent scaffolds: active zone proteins (such as ELKS, Bassoon, RIM, and Munc13) concentrate the exocytic machinery at defined release sites [77], while permanent endocytic scaffolds are constitutively pre-positioned within a confined periactive zone adjacent to active zones [3, 78]. In this architecture, the coupling geometry is physically built into the synapse through permanent scaffolds that persist independently of secretory activity. The global and local coupling we observe in the synchronized constitutive exocytosis, by contrast, operates across a wide, unconstrained plasma membrane surface without obvious permanent infrastructure.

In insulin-stimulated adipocytes, Shape2Fate resolves an inverse coupling regime: productive GSV fusions occur preferentially near pre-existing individual CCPs, enabling rapid capture and immobilization of newly inserted GLUT4 within seconds (Fig. 8C, bottom). These event-resolved measurements support a targeted GSV docking mechanism, and are distinct from the passive diffusion-and-capture model described for VAChT in depolarised PC12 cells [79]. In contrast to the feedback-like coupling observed in synchronized constitutive secretion, the adipocyte data are consistent with a feedforward circuit: adipocytes direct GSV fusion to CCP-rich membrane regions, enabling seconds-scale GLUT4 capture without relying on passive lateral diffusion. Since multiple steps in GLUT4 trafficking - including vesicle tethering, docking, and post-fusion dispersal - are perturbed in insulin-resistant states [39], it is conceivable that the spatial coupling between GSV fusion sites and CCP positioning is itself a point of dysregulation. Shape2Fate could directly interrogate this in metabolic disease models.

Across the two tested systems, Shape2Fate analysed highly dissimilar carrier morphologies (elongated tubules in RUSH; small vesicles in adipocytes) and various CCP distributions (size/density shifts; Fig. 8A, B; Supplementary Fig. 12). That a single geometry-aware framework can resolve carrier trajectories and classify coupling architectures across both contexts motivates the next development steps aimed at more diverse morphologies and trafficking processes. The detector’s generality relies on synthetic training data from an explicit forward model of TIRF-SIM image formation, but this also defines its current boundaries: any carrier morphology not represented in the simulator (e.g., caveolar rosettes, tubular networks, or large static flat lattices) may be missed or misclassified. Extending the simulator’s parametric shape library, guided by EM atlases [80] and published super-resolution data, would directly address this limitation. Where analytical simulation alone is insufficient, generative approaches such as diffusion models or style-transfer domain adaptation can further help to narrow the domain gap between simulated and experimental data [34, 81]. These extensions will be particularly important for processes such as endosomal cargo sorting. Endosomes tubulate, branch, and fission, producing topologies that go beyond the single object representation for which the current distance-map encoding and shape-index descriptor were designed.

As with any microscopy-based approach, Shape2Fate is ultimately bounded by the spatiotemporal resolution of the input data. These bounds manifest in two ways. First, temporally: in neuroendocrine cells and at synapses, exocytic fusion pores can “flicker” on microsecond to millisecond timescales as metastable open/closed intermediates. These transient pore openings allow only partial release of soluble cargo [82, 83]. Because such events are shorter than a single acquisition frame, they cannot be resolved and tracked as distinct states. Shape2Fate may therefore undercount very brief real productive fusions. Second, spatially: vesicles separated by less than the resolution limit may appear as a single object, confounding both detection and trajectory assignment. Our benchmarking analysis directly quantifies this spatial constraint (Fig. 4): the remaining ambiguous CCP tracking cases arose predominantly in dense clusters where overlapping objects could not be optically resolved even by expert human annotators. Together, these temporal and spatial limits define the interpretive scale of Shape2Fate. When ultra-fast or molecular-scale mechanisms are the central question, complementary experimental approaches such as electrophysiology or single-molecule assays remain essential [83].

Looking ahead, improving linking robustness and morphology coverage may broaden Shape2Fate’s capability to other membrane trafficking pathways. The most direct extension of the linking framework is to replace its handcrafted edge costs with learned association functions. Shape2Fate currently scores candidate links by a fixed combination of Euclidean distance and shape-index difference. These costs were calibrated for the shape-evolution profiles of CCPs and exocytic carriers studied here. Transformer-based linkers [23] that learn pairwise associations directly from detection features could absorb pathway-specific dynamics automatically, removing manual tuning as a bottleneck when the framework is applied to a new biological context. An equally promising extension is from two-dimensional TIRF-SIM to volumetric modalities such as lattice-SIM, which would enable tracking of individual carriers from their biogenesis at the Golgi through to fusion. Recent lattice-SIM studies of post-Golgi tubular carriers during constitutive secretion [38, 43] illustrate the kind of morphologically rich data that a volumetric SI and adapted synthetic generator could exploit.

In summary, our results establish that the time-resolved morphology of a single membrane-trafficking carrier, captured by super-resolution live-cell imaging and interpreted by a shape-aware tracker, is sufficient to decode its functional outcome. By exposing, at scale, the spatial rules coordinating exocytosis with endocytosis, Shape2Fate enables mechanistic dissection of the coupling mechanisms across cell types and conditions. The rules uncovered here are not universal but pathway-specific, suggesting that exo-endocytic coordination is a tunable feature adapted to the distinct demands of each trafficking context.

## Methods

### Cell lines and cell culture

Human retinal pigment epithelial (RPE-1 [84]) were obtained from ATCC. SH-SY5Y expressing mEmerald cargo reporter were obtained from Gershlick lab [38]. 3T3-L1 fibroblasts expressing pHluorin-GLUT4 were a kind gift from the Fazakerley lab. RPE-1 and SH-SY5Y and 3T3-L1 cells were cultured in Dulbecco’s modified Eagle’s medium (DMEM) high-glucose GlutaMAX medium (31966-021, Thermo Fisher Scientific-Gibco) supplemented with 10% FBS (Sigma-Aldrich). All cells were cultured under 5% CO2 at 37°C. Cell lines were routinely tested for absence of mycoplasma contamination with MycoAlert Mycoplasma Detection Kit (Lonza).

### Constructs and DNA

Retroviral EGFP-CLCa construct was described previously [85]. SNAP-CLCa, and mRuby2-CLCa were cloned using Gibson assembly in retroviral vector pMIB6 as described earlier [13]. Dynamin2-mRuby3 was obtained from Sandra Schmid lab (UT Southwestern).

### Cell line engineering

The following cell lines were generated using retroviral expression systems in the respective parental cell lines: RPE-1 EGFP-CLCa, RPE-1 SNAP-CLCa, SH-SY5Y SBP-mEmerald-LAMP1 mRuby2-CLCa, 3T3-L1 pHluorin-GLUT4 SNAP-CLCa. Viral particles were produced by co-transfecting HEK 293T cells with a complex formed by polyethylenimine (PEI) and three plasmids for retrovirus production (Gag-Pol, VSV-G, and the retroviral expression construct in a pMIB6 backbone). Cells were sorted post infection using Fluorescence-activated cell sorting (FACS) to isolate populations with low and homogenous expression levels.

### Transient transfection with Dynamin2-mRuby3

RPE-1 SNAP-CLCa cells were transiently transfected with Dynamin2-mRuby3. Cells were seeded onto coverslips in a 6-well plate and allowed to adhere overnight. The following morning, cells were transfected at 50-60% confluence using 2.5ug DNA, 5ul P3000™ Reagent, and 3.75ul Lipofectamine™ 3000 reagent (Thermo Fisher Scientific) in 2.5ml Opti-MEM (Gibco, 31985070) per well. Cells were incubated with the transfection mixture at 37°C and 5% CO2 for 6 hours, before exchanging DMEM supplemented with 10% FBS. Cells were imaged 8 hours post-transfection.

### 3T3-L1 fibroblasts differentiation into adipocytes and stimulation protocol

We adapted the protocol published in [86]. 3T3-L1 pHluorin-GLUT4 and SNAP-CLCa expressing fibroblasts were grown to 100% confluence in a 6-well plate in normal maintenance medium DMEM with high glucose and GlutaMax (Gibco 10566016) and supplemented with 10% FBS (Sigma-Aldrich, F7524). Differentiation was induced 3–4 days after reaching confluency (Day 0) by the addition of 220 nM dexamethasone (Sigma-Aldrich, D4902), 350 nM insulin (Sigma-Aldrich, I9278), 500 µM 3-isobutyl-1-methylxanthine (IBMX; Sigma-Aldrich, I5879) and 410 nM biotin (Sigma-Aldrich, B4639) to the media. On Day 3, cells were changed with post-differentiation media containing 350 nM insulin and cultured for a further 3 days. On Day 6, cells were re-fed with normal maintenance media. For microscopy, coverslips were coated with Matrigel (1:100 in ice-cold media, 30 min at 37°C). Differentiated adipocytes were seeded on Day 6 and imaged between days 8 and 9 with 90% differentiation efficiency confirmed by morphology, lipid-droplet content, and TIRF microscopy (Supplementary Movie 4). On the day of TIRF-SIM microscopy, prior to insulin stimulation, cells were washed three times with warm PBS and incubated for >2 hours in serum-starvation media made of serum-free FluoroBrite DMEM (Gibco, A1896701), 0.2% (w/v) bovine serum albumin (BSA; Bovostar, Bovogen), 2 mM GlutaMAX and 50 mM HEPES. Stimulation was performed with 100 nM insulin added directly to cells. TIRF-SIM Imaging began 5 and 30 minutes post-stimulation.

### Synchronized constitutive secretion with RUSH-mEmerald-LAMP1

SH-SY5Y cells expressing RUSH-mEmerald-LAMP1 were imaged using FluoroBrite™ DMEM (Gibco) supplemented with GlutaMAX™ (Gibco) and 10% FBS (Sigma-Aldrich) before stimulation. For stimulation, 500 µM D-biotin was added directly to the cells [38]. Imaging was done 15 minutes after addition of biotin, which proceeded for a following 30 minutes.

### Transmission electron microscopy

Cells were fixed with 2.5% glutaraldehyde/2% paraformaldehyde in 0.1 M sodium cacodylate buffer, pH 7.2, post-fixed with 1% osmium tetroxide in sodium cacodylate buffer, dehydrated through a graded series of ethanol and embedded in Agar 100 resin. Randomly cut and orientated 60 nm sections were stained with uranyl acetate and lead citrate [87] and images of CCVs were recorded with a 16-megapixel Eagle charge-coupled device (CCD) camera on a Tecnai G2 Spirit BioTWIN transmission electron microscope (FEI, Eindhoven) operated at 80 kV. EM Image analysis and measurement of CCPs was carried out in ImageJ/Fiji [88].

### Live cell TIRF and TIRF-SIM microscopy

Cells were imaged in FluoroBrite™ DMEM (Gibco) supplemented with GlutaMAX™ (Gibco), 10% FBS (Sigma-Aldrich), and 50 mM HEPES unless otherwise stated. All cell lines expressing SNAP-tagged proteins were incubated for at least 1 hour with Janelia Fluor® Dyes (1:1000 dilution) prior to imaging. Cells were washed with media to remove unbound dye immediately prior to imaging. All TIRF imaging was acquired at the CIMR, University of Cambridge, using a Zeiss Elyra PS1 inverted microscope equipped with a 100× 1.49-NA Apo TIRF oil-immersion objective and a PCO Edge 5.5 sCMOS camera. Unless stated otherwise TIRF-SIM experiments were carried out at the Micron Bioimaging Facility, Biochemistry Department, University of Oxford, using a DeltaVision OMX SR inverted super-resolution microscope equipped with a 60× 1.5-NA TIRF UPLAPO oil-immersion objective (Olympus) and a PCO Edge 4.2 sCMOS camera. The microscope was operated using OMX Master Control Software (AcquireSR). Immersion oil matched to the objective specification was used, and the objective correction collar was matched to the acquisition temperature (37°C) to minimize spherical aberration. Excitation was provided by solid-state lasers (488 nm, 561 nm and 639 nm), and the illumination angle was adjusted to achieve total internal reflection at the glass–aqueous interface. Alignment parameters were obtained from calibration measurements with 0.1 µm TetraSpeck microspheres (ThermoFisher; T7279) at the start of each acquisition session.

Imaging was performed in single-plane TIRF-SIM mode. At each time point, nine raw images were acquired in a single focal plane (three illumination orientations × three phase shifts). The effective pixel size at the sample plane was 0.0791 µm. Exposure times were kept as low as possible, typically 10-30ms per raw frame, to maintain signal-to-noise while limiting phototoxicity. For exocytosis experiments, acquisitions were performed at 250 ms per reconstructed frame, whereas endocytosis experiments were acquired at 2–4 s per reconstructed frame depending on experimental design.

For dual-channel endocytosis–exocytosis experiments, the nine raw SIM frames were interleaved between channels (as depicted in Fig. 6D) to ensure that rapid exocytic fusion events were reliably captured. In this configuration, the CME channel was acquired at 8x lower temporal sampling rate than the exocytosis channel.

SIM reconstruction was performed using custom software accessible at https://github.com/harmanea/shape2fate and as described below.

### Channel registration

Multicolour acquisitions were aligned using 0.1 µm TetraSpeck microspheres (ThermoFisher, T7279) as fiducial markers. Bead centroids were localized in each channel and paired across channels, and the corresponding coordinates were used to compute a similarity transformation (scaled rotation) that minimizes the root-mean-square positional error between channels, following the sub-pixel registration approach described in. Multicolour image stacks were then processed using the same transform. The residual RMS registration error, estimated from bead centroids after transformation, was below 30 nm.

### Estimation of Effective Lateral Resolution from TIRF-SIM Images

We analysed the spatial frequency content of TIRF-SIM reconstructions to estimate their effective lateral resolution, following the approach of Ball & Schermelleh [15] (Supplementary Fig. 2). First, we performed thresholding by subtracting the image’s modal intensity to zero the background (SIMcheck utility “Threshold & 16-bit conversion”). The 2D Fourier transform was computed (SIMcheck “Reconstructed data -Fourier plots”) from this background-corrected image. We averaged the Fourier amplitudes in concentric rings (isotropic averaging) to produce a radial frequency profile, which plots integrated signal strength (amplitude) as a function of spatial frequency. Finally, we defined the practical resolution cutoff at the inflection point where the radial profile’s amplitude curve decays to meet this noise floor (marked by an arrow in Supplementary Fig. 2).

### SIM Reconstruction

Nine raw SIM images (three illumination orientations × three phase shifts) are reconstructed in the Fourier domain following established approaches [40, 89]. For each orientation, phase-shifted images are decomposed into zero- and ±1st-order frequency components by solving a linear system. Illumination pattern frequencies are estimated automatically using an optimization-based procedure, optionally initialized with an approximate user-provided estimate (see Parameter estimation).

The separated components are shifted to their correct Fourier positions and combined using a generalized Wiener filter [40]. In this study, the regularization parameter was fixed to w=0.05. To mitigate high-frequency noise amplification, an apodization filter derived from the ideal incoherent optical transfer function (OTF) support is applied (apodization cutoff = 2.0; bend exponent = 0.9) [89].

The OTF is modeled using an exponential approximation of the ideal wide-field OTF [89] with fixed curvature parameter 0.3. Microscope-specific parameters (numerical aperture, emission wavelength, and pixel size) are set according to the acquisition system (typically NA = 1.49–1.50; emission wavelengths 512–527 nm for green, 568–603 nm for red, and 679 nm for far-red channels; pixel size 0.061 µm or 0.0791 µm prior to reconstruction). This model OTF was applied to all tested microscopes and reconstruction tasks.

To improve reconstruction robustness, a constant camera offset (typically 100 intensity units, as defined by the microscope protocol) is subtracted prior to processing, and image borders are tapered using a sin² window over 15 pixels to reduce Fourier edge artefacts. Reconstruction results in exact two-fold upscaling (e.g., 512 × 512 → 1024 × 1024 pixels). All non-microscope-specific reconstruction parameters were fixed across datasets used in this study.

### Parameter estimation

Accurate SIM reconstruction depends critically on the precise knowledge of the illumination pattern parameters: frequency, orientation, phase offset, and modulation amplitude. Illumination parameters can vary between acquisitions and were therefore re-estimated for each dataset. In the case of time-lapse acquisitions, we empirically found it sufficient and more stable to estimate the parameters only at the first time point. Estimation is performed independently for each illumination orientation, using the corresponding triplet of low-resolution raw images.

To improve the robustness and accuracy of parameter estimation, we first pre-process all raw images using Richardson-Lucy deconvolution with 10 iterations with the OTF described above, motivated by the approach of Perez et al. [42]. Deconvolution is applied per frame and used exclusively for parameter estimation.

A coarse estimate of the illumination frequency is obtained either from a previous high-SNR dataset or automatically using the phase-only correlation method of Cao et al. [41]. The automatic estimate is accepted when three equidistant peaks at a consistent frequency are detected; otherwise, an estimate obtained from bead calibration data is used.

Sub-pixel refinement of the illumination frequency also follows the approach of Cao et al. [41]. The coordinates are jointly optimized within ±1.5 pixels of the coarse estimate, with sub-pixel shifts applied via the Fourier shift theorem. The local search is performed by a bounded L-BFGS-B optimisation procedure [90, 91].

After the frequency and orientation have been refined, the phase offset of the pattern is estimated using the estimation-through-reconstruction approach described by Perez et al. [42]. We adapt this method by evaluating the cross-correlation metric at six equally spaced phase offsets between 0 and 2π, fitting a sine curve to the results, and selecting the phase corresponding to the peak of the fitted curve. Six samples were found to be the minimum required for stable fitting while maintaining computational efficiency.

Finally, the modulation amplitude is estimated. In high-SNR cases, this was done using the complex linear regression method described by Gustafsson et al. [40] and later detailed by Müller et al. [89]. However, in low-SNR, live-cell time-lapse data sets, this value could not generally be estimated reliably, so a fixed value of 0.5 was used instead.

### Detection

The goal of this step is to localise individual vesicles and extract information about their shape. Detection is performed independently for each frame of a time-lapse recording. While the overall approach is shared, the detection strategy is adapted slightly for CME and exocytosis to account for differences in appearance and variability of vesicle shapes. We experimented with various network architectures and training settings during development, but the configuration and approach described here were empirically found to perform best on the validation dataset (accessible at https://zenodo.org/records/17484958).

### U-Net neural network architecture

We use a 2D U-Net [58] convolutional neural network with a depth of three levels. At each depth, the spatial resolution is halved, and the number of feature channels is doubled, starting with 16 filters in the first layer. Each block consists of two consecutive 3×3 convolutions (same padding, no bias), each followed by batch normalisation and ReLU activation. Downsampling is performed using 2×2 max pooling, and upsampling is done using nearest-neighbour interpolation followed by a 1×1 convolution to adjust the number of feature channels. A final 1×1 convolution reduces the feature maps to the required number of output channels, followed by a sigmoid activation to constrain the output values between 0 and 1.

The network takes a single-channel input image and produces either a one-channel output (for CME, Fig. 2C, D) or a two-channel output (for exocytosis, Fig. 5B, Supplementary Fig. 5), with separate channels for regular vesicles and fusion events. Input images are normalised per frame to zero mean and unit standard deviation. The network output is a normalized interior-distance map, with high values assigned to the interior of vesicular structures and values smoothly decaying to zero at their boundaries.

### Synthetic Training Data

To train the detection models, we generated synthetic image sequences that mimic TIRF-SIM snapshots. The simulation pipeline replicates both image formation and reconstruction (Fig. 2B; Supplementary Fig. 1).

Each sample begins with the creation of a high-resolution, latent image representing the underlying vesicle appearance. This image is then illuminated with nine structured light patterns, with randomly selected initial orientation and phase offset. Each illuminated image is convolved with a model PSF and downsampled to half the resolution. This results in nine low-resolution images per sample, which are reconstructed using a Wiener filter reconstruction, identical to that used for real data (see SIM Reconstruction).

To improve realism and inference robustness, we apply the following augmentations: foreground and background intensities are randomised, short-distance Brownian motion is introduced across the nine structured illumination images to mimic lateral carrier mobility, background out-of-focus signal, uneven illumination or cell attachment is simulated using Perlin noise [92], and Poisson noise is added to each of the nine raw images before reconstruction, consistent with photon shot noise in fluorescence imaging [93]. Additionally, small perturbations are applied to the illumination pattern parameters to simulate inaccuracies in their estimation.

All simulation parameters can be freely adjusted in the provided image generator (accessible at https://github.com/harmanea/shape2fate). For all presented experiments, we fixed them as follows: Synthetic images were generated at a size of 128×128 pixels, which we found to provide sufficient spatial context for the network while keeping training computationally efficient. The simulated imaging parameters were set to match those of the experimental systems: NA of 1.49, emission wavelength of 512 nm, sensor pixel size of 0.07 µm, Wiener reconstruction parameter of 0.1, and structured illumination frequency of 0.17.

### Synthetic training data for CME

Synthetic CCPs are generated by placing structures onto a predefined 6×6 grid, with grid points spaced 16 pixels from each other and from the image edges. Between 5 and 15 of these positions are randomly selected for vesicle placement. To introduce spatial variability and allow for occasional clustering of CCPs, each selected position is offset randomly by up to ±8 pixels in both x and y directions.

Each CCP is assigned SI, sampled from a beta distribution with parameters a = 1 and b = 2, rescaled to the range [0.1, 1]. The SI determines both the size and morphology of the CCP (Fig. 2A). On a blank high-resolution canvas, CCPs are rendered as near-point-source filled disks at low SI and as annuli at high SI.

For a CCP centred at position c = (x₀, y₀) with maximum radius r₀ = 2.5 pixels (≈87.5 nm), the target radius r = r₀·SI and ring thickness t = 1 pixel, let 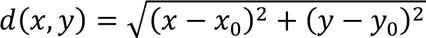. The intensity contribution at pixel (x, y) is given by 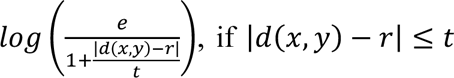, otherwise 0.

The final synthetic image is obtained by summing up the contributions of all vesicles on the canvas. This high-resolution image is then processed through the full TIRF-SIM simulation pipeline (Fig. 2B; Supplementary Fig. 1A).

The corresponding training target is a normalized interior-distance map, where the value at the centre of each vesicle equals its SI and decays smoothly to zero at the boundary (Fig. 2C).

For adipocytes, synthetic data generation was adjusted to account for their larger CCP diameter: CCP radii were scaled up, the SI distribution was shifted so that although the range remained [0.1, 1], the maximum value corresponds to a larger radius r₀ = 3.5 pixels (≈122.5 nm), and a separate model was trained on this adapted dataset.

### Synthetic training data for Exocytosis

Synthetic exocytic vesicles are generated using the same 6×6 grid layout as in the CME case, but due to their larger size, only 2 to 8 positions are selected per sample, and they are randomly offset by only up to ±5 pixels in both spatial dimensions (Supplementary Fig. 1B).

Vesicles are rendered as Bezier curves. For each vesicle, a random line segment (length between 1 and 12 pixels, with shorter lengths appearing as disks) is placed at the selected position and oriented at a random angle. The endpoints of the Bezier curve lie along this segment, while the control point is randomly offset (±5 pixels) perpendicular to the segment to introduce curvature. The intensity of each vesicle is varied randomly.

To balance sensitivity and specificity, a fusion event is added in 30% of training samples. These are simulated by placing 200 point sources per frame, with positions drawn from a normal distribution centred at the vesicle location. The spread of the distribution increases across frames, simulating radial diffusion over time. Each low-resolution image corresponds to a distinct time point in this diffusion process, leading to characteristic reconstruction artefacts that resemble those observed in real data. Fusion events can start before or end after the nine-frame acquisition window, and their intensity is also varied randomly.

To prevent false-positive detections of filopodia, which share morphological features with tubular exocytic carriers, a synthetic filopodium is added to 20% of training samples. It is generated by selecting a random point along the patch boundary and drawing a line towards a random location within the canvas. The filopodium has randomly sampled thickness and intensity, with intensities kept lower than those of the exocytic carriers.

The training target for vesicles is a normalized interior-distance map of the rendered object shape. For fusion events, the target map has a value of 1 at the event centre and decreases smoothly to 0.1 at the radius containing 90% of all simulated points (Fig. 5B).

### Training procedure for CME and exocytosis

Models were trained using a batch size of 16 for 300 training cycles, with 100 batches per cycle. After each cycle, performance was evaluated. Training was performed using the AdamW optimizer [94] with a weight decay of 0.05 and a fixed learning rate of 1.6e-4. The loss function was binary cross-entropy (BCE), computed independently for each pixel and output channel.

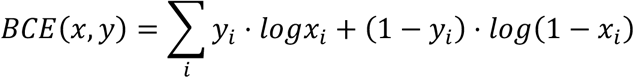

where x_i_ is the output value of the i-th pixel estimated by the network and y_i_ is the corresponding value in the training target.

### Processing of experimental data

When applied to real data, the same per-frame normalisation used during training is applied: each frame is normalised to have zero mean and unit standard deviation. Full-resolution images are passed directly to the trained network.

For CME, candidate CCPs are detected by identifying local maxima in the output distance map that exceed a value of 0.1 (Fig. 2D). This allows the model to resolve clustering CCPs. For each peak, the corresponding SI value is extracted from the map. In rare cases where neighbouring pixels share the same peak value, they are merged into a single detection.

For exocytosis, connected components above a threshold of 0.1 are identified separately in each of the two output channels. Detections are classified as vesicles (SI=0) or fusion events (SI=1) based on which channel they appear in (Fig. 5B, Supplementary Fig. 5). Empirically, we found that retaining only components containing values above 0.4 (vesicles) or 0.5 (fusion events) effectively suppresses spurious false positive detections.

### Linking

The goal of the linking step is to connect vesicle detections across frames to form complete trajectories. Our approach performs this step globally across the entire time-lapse sequence, rather than relying on frame-by-frame association. It is based on a previously published method by Löffler et. al [52], but we introduce several important modifications and therefore describe it in more detail here.

The linking process is divided into two stages (Fig. 2E). First, an initial matching step links detections over time into tracks, allowing for both splits and merges. This flexibility helps compensate for false detections (false positives) or missed detections (false negatives). In the second stage, an untangling step selects suitable track edits to resolve these branching events. Both stages are formulated as integer linear programming (ILP) problems and solved using the SCIP solver [95].

### Matching step

To construct vesicle trajectories from detections over time, we formulate the matching problem as a global coupled minimum-cost flow optimisation. The objective is to compute a flow that assigns each detection to a biologically plausible trajectory, minimizing the total cost of selected transitions and yielding a set of disjoint paths through the graph that represent candidate vesicle tracks. We model the full time-lapse sequence as a directed acyclic graph G = (V, E), where:

- **Nodes V**:
- `o` Flow source (q^-^) and sink (q^+^)
- `o` Detection nodes o_i,t_ for each detection i at time t
- `o` Auxiliary nodes for:
- ▪ Appearances a_t_, disappearances d_t_
- ▪ Skips x_i,t’_ (for t’ > t)
- ▪ Splits s_i,t_ and merges m_i,t_
- **Edges E** connect nodes between and within frames to model possible transitions:
- `o` q^-^ → a_t_, o_i,0_
- `o` o_i,t_ → o_j,t+1_, x_i,t’_, d_t+1_, s_i,t+1_, m_k,t_; o_i,T_ → q^+^
- `o` s_i,t_ → o_j,t_
- `o` m_i,t_ → o_i,t+1_, d_t+1_
- `o` x_i,t_ → o_j,t+1_, m_k,t_
- `o` a_t_ → d_t+1_, o_i,t+1_, s_j,t+1_
- `o` d_t_ → q^+^

We formulate an ILP with integer flow variables z^f^(u, v) ∈ ℕ_0_ and costs c(u,v) ∈ ℝ for each edge (u, v) ∈ E, minimizing total cost:

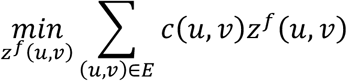

subject to:

- **Flow conservation:**

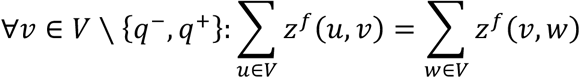
- **Detection assignment:**

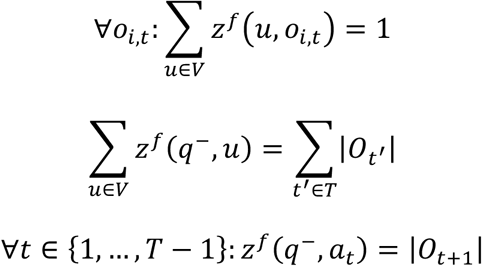

where T is the set of all time points in the graph and |O_t_| is the number of detected objects at time point t.

- **Capacity constraints:**
- Each edge (u, v) is assigned a maximum capacity b(u, v) which restricts the flow over that edge:

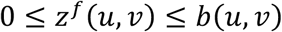
- Most edges have a capacity of 1, except for the following:

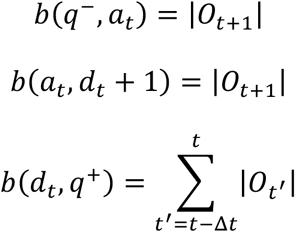

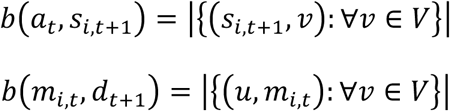

where Δt is the maximum number of skipped time points.
- **Merge constraints:**

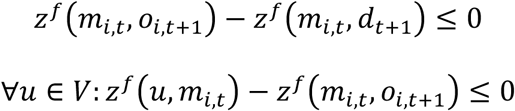
- **Split constraints:**

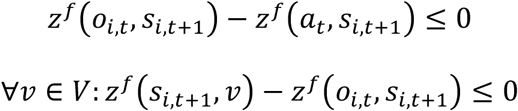

Edge costs c_e_ are set as follows:

- Most costs are set to 0.
- The cost of edges between detections c(o_i,t_, o_j,t+1_) is based on their Euclidean distance and shape feature differences.
- The cost of skip edges is c(x_i,t’_, o_j,t’’_) = c(o_i,t_, o_j,t’’_) · (t’’-t).
- The cost of merge edges is c(o_i,t_, m_j,t_) = c(o_i,t_, o_j,t+1_).
- The cost of split edges is c(s_i,t_, o_j,t_) = c(o_i,t-1_, o_j,t_).
- The cost of appearance edges c(a_t_, o_i,t+1_) and disappearance edges c(o_i,t_, d_t+1_) are a hyperparameter.

To decrease the size of the graph and accelerate optimisation, candidate edges between detection nodes are retained only if their cost is below a given threshold hyperparameter. Split and merge nodes are considered only for the edges remaining after this pruning. Skip edges are only created between detections whose cost would be below the same threshold if they were in the neighbouring frames.

For CME, the cost of edges between detections is defined as the sum of their Euclidean distance and the square difference of their SIs. The inclusion of the SI term was assessed in an ablation study, omitting it resulted in reduced association performance (Supplementary Table 1). The cost of appearance and disappearance is set to a constant of 5. The maximum pruning distance is set to 7.5.

For exocytosis, the cost of edges between detections is defined as their Euclidean distance. The cost of appearance and disappearance is set to a constant of 10. The maximum pruning distance is set to 15.

Optimal hyperparameters were found experimentally by a parameter sweep on the respective validation tracking datasets.

After the optimisation problem is solved, the nonzero flow values define a set of paths through the graph, each corresponding to a sequence of linked detections. These paths are extracted as preliminary trajectory segments, which we refer to as *tracklets*. At this stage, tracklets may still contain branching events due to splits or merges. These ambiguities are resolved in the subsequent untangling step.

### Untangling step

To resolve complex tracklet branching, we construct their graph with edges connecting splitting and merging tracklets and define a set of possible operations with defined costs to alter their structure. The goal is to select a subset of the operations with the minimum cost that output a set of tracks without any splits or merges.

### Operations

- Edge removal z^e^_ps_ between a tracklet ω_p_ and ω_s_ (boolean)
- `o` cost = *γ*, where *γ* is a hyperparameter, typically set to 10
- Split tracklet ω into k parts z^s^_n_ (integer)
- `o` cost = k · |ω|, where |ω| is the length of tracklet ω
- `o` When a tracklet is split, k new tracklets are created instead of it. If the original tracklet had both predecessors and successors, they are connected via the newly created tracklet based on their Euclidean distance. The remaining predecessors or successors are connected to the remaining new tracklets. The detection positions of the new tracklet are averaged with the last position of the predecessor and the first position of the successor.
- Merge k tracklets into one z^m^_r_ (boolean), where r is a multi-index of a set of tracklets
- 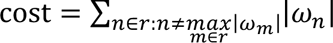

e.g. the sum of the lengths of all the merging tracklets except the longest one

- A set of 2 or more tracklets can only be merged if they:

▪ share the same predecessors and successors (one of which may be empty)

▪ share the same successors and some tracks have no predecessors but start later or at the same time as the track(s) with a predecessor

▪ share the same predecessors and some tracks have no successors but end earlier or at the same time as the track(s) with a successor

- If an edge is cut from a tracklet in a merging set, the newly created merged tracklet will be disconnected from the other tracklet even if it was connected to a different tracklet in the merging set. Moreover, it is enforced via a constraint that only a single such edge may be cut between a tracklet and a merging set. This might be a limitation in very rare cases, but it is necessary to correctly define the operation constraints.

### Constraints

- Each tracklet has at most one predecessor and one successor after all operations are applied
- `o` Predecessor constraint for tracklet ω_n_:

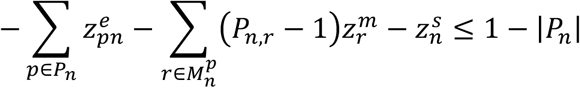

where P_n_ is a set of predecessors of tracklet ω_n_, M^p^_n_ is a set of all sets of tracklets that can be merged that contain predecessors of tracklet ω_n_ and P_n,r_ is the number of tracklets from the set of mergeable tracklets r, that are a predecessor of ω_n_.
- Successor constraint for tracklet ω_n_:

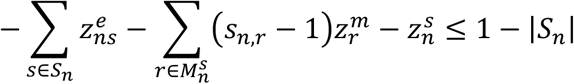

where S_n_ is a set of successors of tracklet ω_n_, M^s^_n_ is a set of all sets of tracklets that can be merged that contain a successor of tracklet ω_n_ and S_n,r_ is the number of tracklets from the set of mergeable tracklets r that are a successor of ω_n_.
- The intuition behind these constraints is that for each extra predecessor or successor, one of the operations on the left has to be performed.
- A tracklet can be merged with at most one set of tracklets, or it can be split

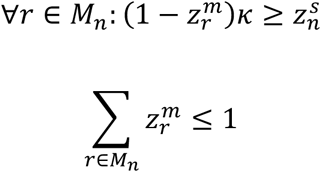

where M_n_ is a set of all sets of tracklets that can be merged with tracklet ω_n_ and *k* is a large constant (e.g. equal to the total number of tracklets)
- At most one edge to a merging set r of tracklets may be removed

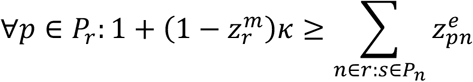

where P_r_ is a set of all predecessors of all tracklets in the set of mergeable tracklets r.

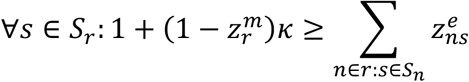

where S_r_ is a set of all successors of all tracklets in the set of mergeable tracklets r.

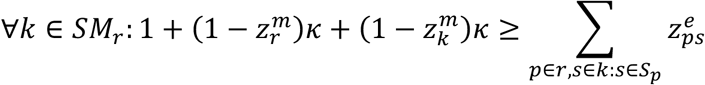

where SM_r_ is a set of all tracklet merge sets that contain a successor of any of the tracklets in r.

The ILP selects a subset of operations minimizing total cost while enforcing the above constraints.

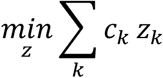

After solving the untangling optimisation, the selected operations are applied to the tracklet graph. The updated graph is then traversed to connect sequential tracklets, each now with at most one predecessor and one successor, into final trajectories. These complete tracks represent consistent, unbranching sequences of detections and form the final vesicle trajectories used for downstream analysis.

### Filtering

Trajectories shorter than the minimum required length are discarded, as they typically arise from spurious detections and are not informative for downstream analysis. In this study, we used a minimum length of five frames. For exocytosis data, trajectories in which the SI equals 1 for more than 75% of their duration are also removed, as these predominantly correspond to noise rather than true events. For adipocyte exocytosis experiments, this filtering was not applied, because many bona fide fusion events produced very short productive trajectories (often shorter than five frames) in which the majority of detections had SI = 1.

### Productivity classification

Following trajectory extraction, each track is classified as either productive or non-productive. The criteria differ between CME and exocytosis to reflect the distinct morphological and temporal features of each process.

### Productivity classification for CME

For CME, a trajectory is classified as productive if at least one detection within the track reaches SI greater than 0.7. This threshold was determined by maximizing agreement (F1 score) with manual productivity annotations on the validation dataset.

For adipocytes, CCPs are larger and synthetic training data were generated with an increased maximum radius (see section *Synthetic training data for CME*), resulting in a shifted SI distribution. The productivity threshold was therefore re-estimated on the validation data, yielding an optimal value of 0.45.

### Productivity classification for Exocytosis

For exocytosis, trajectories are first preselected as potentially productive if they contain at least one detection classified as a fusion event within the final five frames of that trajectory’s lifetime. These candidates are then evaluated by a dedicated neural network classifier to determine their productivity (Fig. 5C).

The classifier takes as input 45 original low-resolution TIRF frames (5 timepoints × 9 SIM frames), each represented as a 21×21 pixel patch centred on the detection. The model begins with a convolutional stem consisting of four 3×3 convolutional layers (same padding, no bias), each followed by batch normalization and ReLU activation. The first two layers use 8 filters, followed by a 3×3 max pooling layer that reduces the spatial resolution to 7×7, with the final two convolutional layers expanding the representation to 24 channels. Global max pooling is then applied to remove the spatial dimension resulting in a sequence of 45 tokens of dimension 24. Sinusoidal positional embeddings are added to this sequence, which is then passed through a dropout layer with a probability of 0.1. A transformer encoder with three layers, each with four attention heads and a 48-dimensional feedforward layer, processes the embedded sequence. The output is averaged across timepoints, passed through a second dropout layer with a rate of 0.5, and finally fed into a linear layer with sigmoid activation to produce the productivity prediction.

The model was trained on a human-annotated dataset comprising 4,192 trajectories from 34 cells (18 adipocyte, 16 RUSH) and validated on 1,571 trajectories from 12 independent cells (5 adipocyte, 7 RUSH). Class distributions were: adipocytes — train 2,532 negative / 712 positive, validation 719 / 183; RUSH — train 422 / 526, validation 346 / 323. Training used a batch size of 8 for 300 epochs with the AdamW optimizer (weight decay 0.05, fixed learning rate 1e-4), and BCE loss. The model achieved 93% accuracy on the validation set (Supplementary Table 4).

### Computational performance

All computational benchmarks were performed on the CME validation dataset (120 frames, 1024×1024 pixels) using a workstation equipped with an AMD Ryzen 9 5900X CPU and an NVIDIA GeForce RTX 3060 Ti GPU. The table below summarizes the runtimes of each processing stage. All results report wall-clock times averaged over five runs with the first warm-up run discarded.

Reconstruction and detection time complexity scales linearly with the number of frames because each frame is processed independently. Both steps are accelerated substantially by running the core operations on the GPU, which can execute the underlying Fourier transforms, convolutions, and per-pixel operations in massively parallel fashion.

Linking and untangling runtimes depend mainly on the density of objects in the field of view, not only on the number of frames. Linking is computationally demanding because graph building must create all detection nodes, auxiliary nodes, and the edges between them. The solution-finding step is the slowest because it must evaluate the full spatiotemporal context. Solution extraction is fast as it only selects the chosen linking edges. Untangling is simpler in graph construction and solving because only edges between tracklets are considered. However, its solution-extraction step is more expensive since it must apply all operations, including edge removal, tracklet splitting, and merging.

| Stage | Processing step | CPU time (s) | GPU time (s) |
| --- | --- | --- | --- |
| **Reconstruction** | Preprocessing | 1.09 | — |
|  | Parameter estimation | 6.77 | — |
|  | Reconstruction | 97.73<br>(0.81 per frame) | 3.53<br>(0.03 per frame) |
| **Detection** | Detection | 47.84<br>(0.40 per frame) | 6.58<br>(0.05 per frame) |
| **Linking** | Graph building | 38.94 | — |
|  | Solution finding | 88.50 | — |
|  | Solution extraction | 0.64 | — |
| **Untangling** | Graph building | 0.33 | — |
|  | Solution finding | 0.15 | — |
|  | Solution extraction | 20.92 | — |

### Analysis Masking

Each timelapse movie was manually inspected, and typically a single mask was drawn by an expert to include only in-focus and well-adherent cells. If the cell changed shape over time, the mask was designed to remain valid throughout the entire sequence. In the few cases where cell shape changed substantially, multiple masks per movie were used. To prevent distortions caused by edge tapering and to ensure that tracked objects do not move out of view, the outermost 15 pixels on all sides of the field of view were also masked out. Movies exhibiting strong photo-bleaching or photo-toxicity at later timepoints were trimmed. Only trajectories fully contained within the masked region were included in the analysis.

### Normalization of fluorescence intensities across samples and data sets

To compensate for bleaching and enable comparison across different samples, intensity normalisation was applied independently to each timelapse [96]. For each frame, the mean intensity within the masked region was measured and a bi-exponential function was fitted to the resulting time series:

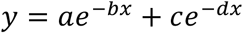

where x is time, and a, b, c, d are fit parameters. The raw intensities were divided by the fitted curve, ensuring that the average intensity within the masked area is 1 across all frames and samples.

For normalisation of clathrin signal, the procedure was modified to normalise only the background. Specifically, the mean intensity was computed using only pixels with values below 0.1 in the U-Net detection output (i.e., excluding CCPs). This ensures that the background intensity, rather than the entire cell, is normalised to 1.

### Generation of TIRF equivalents from TIRF-SIM data

To generate conventional TIRF images from TIRF-SIM acquisitions, the nine low-resolution raw frames corresponding to a single timepoint (with varying structured illumination patterns) were averaged. This suppresses the structured illumination modulation and approximates a conventional TIRF acquisition.

### Sampling of fluorescence intensities

Fluorescence intensities were extracted after per-frame normalization at TIRF resolution (see section *Generation of TIRF equivalents from TIRF-SIM data*), thereby avoiding potential biases introduced by SIM reconstruction. For each detection, its coordinates were downsampled to the TIRF resolution, rounded to integer pixel positions, and the fluorescence intensity was computed as the mean over a 5×5 pixel neighbourhood centred at that location. The neighbourhood averaging reduces pixel-level noise and provides a robust local intensity estimate.

### CCP cluster size

To quantify spatial grouping of visually touching or overlapping CCPs, the U-Net distance map is thresholded (values > 0.1) and partitioned into connected components. The threshold of 0.1 corresponds to the lower bound of the synthetic training target distance transform and so values below this level are not expected to represent valid CCP interiors. Individual CCP detections (see section *Processing of experimental data*) are assigned to the component label at their rounded pixel location, and cluster size is defined as the number of detections within the same connected component.

### Experimental validation of the SI-based productivity criterion using Dynamin2-mRuby3 recruitment

To validate the SI-based productivity threshold, we analysed dynamin recruitment dynamics in CCP trajectories that were classified as productive (see section *Productivity classification for CME*), fully contained within the duration of the movie (i.e. initiated after the first frame and completed before the last), and not part of any cluster (to avoid contamination from nearby CCPs, see section *CCP cluster size*).

Because CCP lifetimes vary, trajectories were temporally normalized. For each trajectory, measurements were linearly interpolated and resampled to 100 equally spaced timepoints spanning its lifetime (normalized time 0–1). This resolution was chosen to preserve temporal detail while maintaining computational efficiency, as typical CCP lifetimes were substantially shorter than 100 frames (200s).

Dynamin fluorescence intensities were normalized per sample and extracted at each resampled detection position using the TIRF-resolution sampling procedure described above.

To assess recruitment dynamics before and after the tracked CCP lifetime, the analysis window was symmetrically extended by 25% of the trajectory duration in both directions, provided that the extended window remained within the movie bounds. During extension, dynamin signal was sampled at the last known CCP position.

Aggregated traces are shown as mean ± SEM across trajectories (Fig. 3B). For visualization, mean traces were smoothed using a normalized moving-average filter of size 5.

For the assessment of maximum clathrin intensities as a productivity criterion (Fig. 3C), clathrin fluorescence intensities for each trajectory were extracted using the same procedure described in [18].

### Manual data annotation for CME and exocytosis

Ground-truth annotations were generated using a custom tool written in MatLab [13] (accessible at https://github.com/Ka-me-nik/TIRF-SIM). This tool provides a graphical user interface that allows the users to add, remove, split, merge, extend or shorten trajectories as well as to add custom tags to individual frames or to adjust the position of each detection.

Annotations were performed on these datasets accessible at Zenodo at https://tinyurl.com/shape2fate: “CME tracking validation”, “CME tracking testing”, “Exocytosis tracking validation”, “Adipocytes-CME coupling”, and “RUSH-CME local coupling”.

For the “CME tracking validation” dataset, the same region was annotated independently by three expert annotators. For the “CME tracking testing” and “Exocytosis tracking validation” datasets, annotation was performed by a single expert. For the “Adipocytes-CME coupling” and “RUSH-CME local coupling” datasets (used for productivity classification), two experts annotated trajectories independently without data overlap.

Initial trajectories were generated by the current version of Shape2Fate and provided as editable starting points. Annotators were instructed to review and correct all trajectories within a designated cell region across the full movie duration, including adding missing detections and modifying or removing faulty ones. Model predictions served only as an initial estimate, final annotations reflect manual expert judgement. Annotators were free to add, remove, split, merge, extend, shorten, and reposition detections and trajectories as needed. For CME, annotators also identified and labelled frames in which CCPs displayed an annular morphology, corresponding to high SI values. All annotations were performed exclusively in the CLCa channel, no additional information or molecular markers were available to or used by annotators.

For exocytosis, annotators marked the frame(s) of cargo release, identified by the characteristic signal diffusion, as the ground-truth label for productive events.

In a separate task for the “Adipocytes-CME coupling” and “RUSH-CME local coupling” datasets, annotators reviewed preselected candidate fusion events and classified them as productive or false positives, with the option to add additional events. No trajectory tracking was performed in this task.

### Tracking metrics

We calculate MOTA and MOTP as described in [63]. For HOTA [64], we adapt the localization similarity measure to our setting, which uses point detections rather than bounding boxes or segmentation masks. Specifically, we use the clipped inverted normalised distance, defined as 1 – min(1, distance/max_distance), which, like IoU, ranges from 0 (no match) to 1 (perfect match). We evaluate HOTA using a single localization threshold α = 0, meaning all distances below the maximum distance are considered for matching. We also report HOTA’s detection accuracy (DetA) and association accuracy (AssA) components.

To better reflect biological requirements such as full trajectory retention for lifetime analysis, we introduce a custom metric somewhat similar to Identification F1 score (IDF1) [97]: the mean temporal intersection over union (µTIOU). The underlying temporal intersection over union (TIOU) is defined as the number of frames where both trajectories contain detections within the matching distance, divided by the number of frames where either trajectory has any detection. This value ranges from 0 to 1, with 1 indicating perfect temporal overlap. If no valid overlap exists, a score of *-infinity* is assigned. µTIOU is calculated by computing TIOU between all track pairs from two track sets, identifying the bipartite matching that maximizes the overall TIOU sum, and then calculate the final score as twice the sum of TIOUs for the matched trajectories divided by the total number of trajectories in both sets.

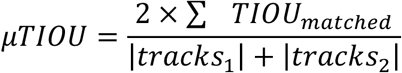

For all metrics, the maximum matching distance is fixed at 5 pixels (≈175 nm) for CME, which is equal to the mean diameter of a CCP, and at 10 pixels (≈350 nm) for exocytosis, to account for the larger carrier size.

### CME tracking datasets for cross-platform evaluation

To evaluate robustness across acquisition platforms, CME tracking performance was assessed on two distinct TIRF-SIM datasets: the publicly available “CME tracking validation” dataset and the independently acquired “CME tracking testing” dataset (see Data Availability).

The validation dataset was previously published and acquired on a custom-built TIRF-SIM setup (100x magnification, 1.49 NA) [13].

A dedicated testing dataset was acquired for this comparison at the Nikon BioImaging Lab, Leiden, on a commercially available Nikon N-SIM S microscope equipped with a CFI SR HP Apochromat TIRF 100× oil-immersion objective (NA 1.49) and an ORCA Fusion BT sCMOS camera (Hamamatsu). The dataset, originally acquired at 0.194 s/frame, was temporally downsampled to ∼2 s/frame prior to evaluation to match standard acquisition conditions for quantitative evaluation. To avoid sampling bias, performance was evaluated across all possible disjoint frame subsets drawn at each frame rate, with tracking statistics averaged across subsets.

### Generation of CCP trajectories with cmeAnalysis and TraCKer for comparison

To evaluate their performance, cmeAnalysis (v. 2024) [18] and TraCKer (v. 2025) [19] were run on TIRF sequences simulated from annotated TIRF-SIM data (see section *Generation of TIRF equivalents from TIRF-SIM data*) (Fig. 4A,B). The detection positions were then upsampled to match the original resolution. Where possible, default or automatically computed values were used. Manual input parameters were optimized to achieve the lowest sum of ranks across the tracking metrics (see section *Tracking metrics*) on the validation dataset, and these were then reused on the testing dataset without any further tuning (Supplementary Tables 1-2). For TraCKer we used the default window size of 5 and the filter threshold of 20, which we found to be optimal. For cmeAnalysis, we used the PSF derived from the physical model. Since cmeAnalysis may return compound tracks containing splits and merges, which cannot be directly compared against linear ground-truth trajectories, we tested several post-processing steps to decompose them into linear segments prior to evaluation. This evaluation revealed that retaining all the individual tracks from the compound was the optimal approach and used this for both the validation and testing datasets.

### Blind paired comparison of Shape2Fate and human-annotated CCP trajectories

To evaluate the quality of trajectory reconstructions, we conducted a blind paired preference test with three independent human reviewers (distinct from the three annotators who generated reference trajectories, Fig. 4C). Each reviewer was presented with short movie cutouts centred on a trajectory mismatch region, spanning ±10 frames (21 total) and ±20 pixels (41 total) around the region of interest (ROI). Within each cutout, two sets of trajectories (labelled by different human annotators or tracked automatically by Shape2Fate) were overlaid in different colours. Reviewers were instructed to select the trajectory set that more faithfully represented the underlying vesicle dynamics visible in the image data, or to abstain if undecided. Either one trajectory from one of the sets, or one trajectory from both sets, was highlighted, and reviewers were instructed to base their decision primarily on this trajectory within ±2 frames of the ROI, while still considering the surrounding temporal and spatial context.

In total, 285 trajectory pairs were evaluated. Of these, 210 comparisons involved Shape2Fate versus a human annotator (70 against each annotator), while the remaining 75 pairs compared trajectories between different annotators to eliminate potential reviewer bias. To avoid systematic effects, the colour assignment of trajectory sets, and the order of ROI presentation were randomized.

ROIs were selected based on mismatches between trajectory sets. To guide this selection, a complexity map was constructed by accumulating all detection and identity mismatches (as defined by the MOTA metric) between the three annotators into a 3D array, where each mismatch contributed a value of 1 at its corresponding space–time location. This map was then smoothed with a Gaussian kernel (σ = 3 px). For each comparison pair (Shape2Fate vs. annotator or annotator vs. annotator), mismatches were identified separately, and those occurring too close to the temporal boundaries of the movie were discarded. A subset of the remaining mismatches was then sampled with probability proportional to the complexity map value at that location, biasing selection toward regions with higher annotator disagreement. To avoid overlap, once an ROI was selected, no additional ROI was allowed within 10 frames or 10 pixels of it.

Reviewer consensus was assessed on a strict unanimity basis. For each trajectory pair, only cases in which all three reviewers independently selected the same method (either Shape2Fate or human annotation) were considered a decisive outcome. If at least one reviewer disagreed, or was unable to choose, the comparison was classified as *undecided*.

### Quantification of clathrin signal explained by CCP detections

The amount of clathrin signal that cannot be explained by CCP detections is calculated by taking the original image and subtracting from it a synthetically generated image that was generated from these detections. For this to work, several important details must be noted:

1. The effect of reconstruction is removed by working on the simulated TIRF version of the original high-resolution image (see section *Generation of TIRF equivalents from TIRF-SIM data*). Similarly, the synthetic image is generated at the lower TIRF resolution.
2. As the cell background is not part of the synthetic image generation process and is irrelevant to this task, it is first subtracted for each frame. It is estimated using a median filter with a circular kernel with a radius of 13 pixels. Negative values after subtraction were clipped to zero.
3. For various reasons, CCPs in the same lifetime phase may have different intensities. To compensate for this, each CCP is given a different weighting in the synthetic image. These factors are determined using least squares fit to minimize the absolute difference between the original image and a synthetic image generated with these weights. As the intensity of nearby CCPs may influence the outcome, all weights are optimized simultaneously. However, this can be very computationally intensive, as a new, large synthetic image must be generated for each iteration of the optimisation process. To speed up the process, the synthetic image is generated on the GPU, and Jacobian values in the fitting algorithm are limited to the immediate vicinity of each CCP.

Once the final difference between the original and synthetic images is obtained, it is filtered using a 15×15 pixel Gaussian filter with a sigma of 2, to limit the effect of slight localisation errors and noise. The result is then clipped from below to include only the significant signal missing from the synthetic image. Through empirical testing, we found that a good clipping threshold is the median of all intensity weight factors divided by 100.

This map of residuals can be visually inspected (Supplementary Fig. 4) or summed to obtain the total amount of unexplained signal. When this is compared to the total signal in the synthetic image, the ratio of the overall signal explained by the CCP detections in that image is obtained.

### Endocytosis-exocytosis RUSH coupling methodology

CCP initiations are considered for trajectories that occur within the cell mask and at least 30 seconds before the end of the movie, to determine the productivity of that track. The CCP initiation density in each spatiotemporal window is calculated by dividing the number of CCP initiations inside the window by its volume (Fig. 6).

The CCP initiation density ratio is calculated by dividing the CCP initiation density in each spatiotemporal region of a cell by the CCP initiation density of the rest of the cell. As the global CCP initiation density varies throughout the duration of the movie and between different cells, the weight of each CCP initiation is inversely proportional to the global initiation density at that timepoint. Additionally, because the global initiation density estimate can be noisy, it is first smoothed using a uniform filter over 10 frames (∼30 s). The considered region is the union of spatiotemporal windows centred on the locations of exocytic fusion sites. These windows are parametrized by their spatial radius, temporal size, and temporal offset (i.e. how many frames before or after the exocytic event they are centred, Fig. 6A,B). These parameters were selected by maximizing the mean productive CCP initiation density ratio in an initial subset of 13 manually reviewed RUSH SH-SY5Y cells from the “RUSH-CME local coupling” dataset (Fig. 6C). This subset represented the fully annotated data available at the time of parameter selection and was used exclusively for window-parameter tuning. The final analysis was subsequently applied to the full dataset without further adjustment.

Despite the large number of events, CCP initiation density maps are relatively sparse. To mitigate this issue when visualising 2D plots of CCP initiation densities, each initiation is expanded to a spatiotemporal region with a radius of 3.5 pixels and seven frames. To eliminate orientation bias, all aggregated frames are randomly rotated first (Fig. 6E,F).

Prior to computation, channels are registered, positions are rounded to integer values, and frame rates are aligned.

### CCP productivity ratio

The CCP productivity ratio was calculated as the number of productive trajectories (*see Productivity classification for CME*) divided by the total number of trajectories, considering only those fully contained within the movie duration (Supplementary Fig. 7).

### CCP size measurement

The radius of each CCP detection is estimated by fitting a Gaussian function to the radial clathrin intensity profile. The profile is computed by assigning y-coordinates as the normalised intensity values (clathrin intensity minus 1, such that the mean background is zero) and x-coordinates as the radial distance from the detection centre. Distances greater than 10 pixels are excluded. The detection centre is defined as the centroid of the connected component where the detection network’s distance map output exceeds 0.1. The final CCP radius is taken as the mean (μ) of the fitted Gaussian.

To quantify radial symmetry, a symmetry score is calculated from the polar transform of clathrin intensity values within a 7-pixel radius of the centroid. The angular profile is computed by averaging across radial coordinates in polar space. For each pixel, the difference from the angular profile at the corresponding radius (linearly interpolated) is computed. The symmetry score is defined as one minus the mean of the absolute differences. Intensity values are normalised by subtracting 1 and dividing by the amplitude of the Gaussian fit.

CCP size evaluation is restricted to productive trajectories that are fully contained within the movie duration and are not part of any cluster. Only frames that meet the following criteria are considered: successful Gaussian fit, amplitude (A) > 1, mean (μ) > 1.25 pixels, standard deviation (σ) < 1.5, symmetry score > 0.75, and SI > 0.7. These thresholds were used to exclude failed or low-confidence fits (e.g., near-flat profiles, very sharp peaks, or overly broad profiles inconsistent with a ring-like CCP footprint) rather than to tune the measured sizes. Among these, the frame with the largest Gaussian mean (μ) is selected as the representative CCP size of the given productive trajectory (Fig. 7A).

### Relative CCP density

CCP density is calculated in a similar way to CCP initiation density, but for all detections along the CCP trajectory. Each detection is weighted inversely proportionally to the smoothed global CCP density at its timepoint, ensuring that the mean global CCP density is 1 at all timepoints and in all cells.

As with initiation density, 2D plots of CCP densities are visualised by expanding each initiation to a region with a radius of 3.5 pixels and randomly rotating all aggregated frames (Fig. 7D, E).

### Normalized CCP proximity of exocytic fusions

To allow comparison of cells with different CCP densities, the relative mean distance of exocytic fusion events to CCPs is computed independently for each cell (Fig. 7F). The value was defined as the ratio of the mean distance between exocytic fusion events and their nearest CCP to the mean distance between randomly chosen locations and their nearest CCP. Random locations were sampled uniformly within the cell mask, with the number of samples matched to the number of observed fusion events in the given cell.

A value of 1 indicates that the positions of exocytic fusion events are random relative to CCPs; a value above 1 indicates that exocytic fusion events avoid CCPs; and a value below 1 indicates that they tend to occur close to CCPs.

### Statistical analysis

Temporal profiles (e.g., normalized fluorescence intensities or SIs) are shown as mean ± standard error of the mean (SEM). Scalar distributions are visualized using boxplots, where boxes span the first to third quartile (Q1–Q3), the horizontal line indicates the median, whiskers extend to 1.5 times the interquartile range, and outliers beyond the whiskers are shown as individual points.

To compare a single observed distribution to an expected mean, a one-sample Student’s t-test was used. Comparisons between two distributions were performed using Welch’s t-test, which is robust to unequal variances and moderate deviations from normality. All statistical tests were two-sided unless stated otherwise.

Sample sizes (n) and p-values are reported in the figure captions or in the main text where relevant. Differences were considered statistically significant when p < 0.05.

## Code availability

The Shape2Fate pipeline is released as an open-source Python package on GitHub (https://github.com/harmanea/shape2fate), which includes example scripts and is the recommended interface for batch or large-scale processing. Google Colab notebooks are provided for single-video reconstruction, detection, linking, and basic analysis of endocytic and exocytic events, and also include demonstrations of synthetic data generation and model training. They are designed for a broad biological audience without programming experience and include both example datasets and support for user-supplied movies.

The graphical annotation tool used for manual trajectory curation is also available as open-source software on GitHub (https://github.com/Ka-me-nik/TIRF-SIM).

All code, notebooks, datasets, and documentation are accessible through the project website (https://shape2fate.utia.cas.cz/).

## Data availability

All raw TIRF-SIM datasets used for analysis and validation, including human annotations, are publicly available on Zenodo (https://zenodo.org/records/17484958)

## Funding

Dr Zuzana Kadlecova: Sir Henry Dale Fellowship of the Wellcome Trust - 220597/Z/20/Z, GAČR 21-16786M, GAČR25-15933S

Prof. Filip Sroubek: GAČR25-15933S Adam Harmanec: GA UK No. 104223

Dr Daniel Fazakerley: MRC Career Development award (MR/S007091/1) and a Project grant (MR/Z504592/1).

Dr David C. Gershlick: Sir Henry Dale Fellowship from the Wellcome Trust (210481) and a Biotechnology and Biological Sciences Research Council (BBSRC) Responsive Mode Grant (BB/W005905/1).

## Acknowledgment

The authors gratefully acknowledge Dr Deirdre Kavanagh and Dr Niloufer Irani at the Micron Bioimaging Facility, Department of Biochemistry, University of Oxford, for their help and enthusiasm for live-cell super-resolution imaging. We thank Dr Reiner Schulte and Gabriela Grondys-Kotarba of the CIMR Flow Cytometry Core Facility, Cambridge Institute for Medical Research, University of Cambridge, for their assistance with cell sorting. We are sincerely grateful to Jérôme Boulanger (MRC Laboratory of Molecular Biology), Philippe Roudot (Institut Fresnel), Assaf Zaritsky (Ben-Gurion University of the Negev), and Professor Michal Kozubek (Centre for Biomedical Image Analysis, Faculty of Informatics, Masaryk University) for their advice and thoughtful suggestions and careful critical reading of the manuscript.

## Supplementary information

**Supplementary Figure 1.**
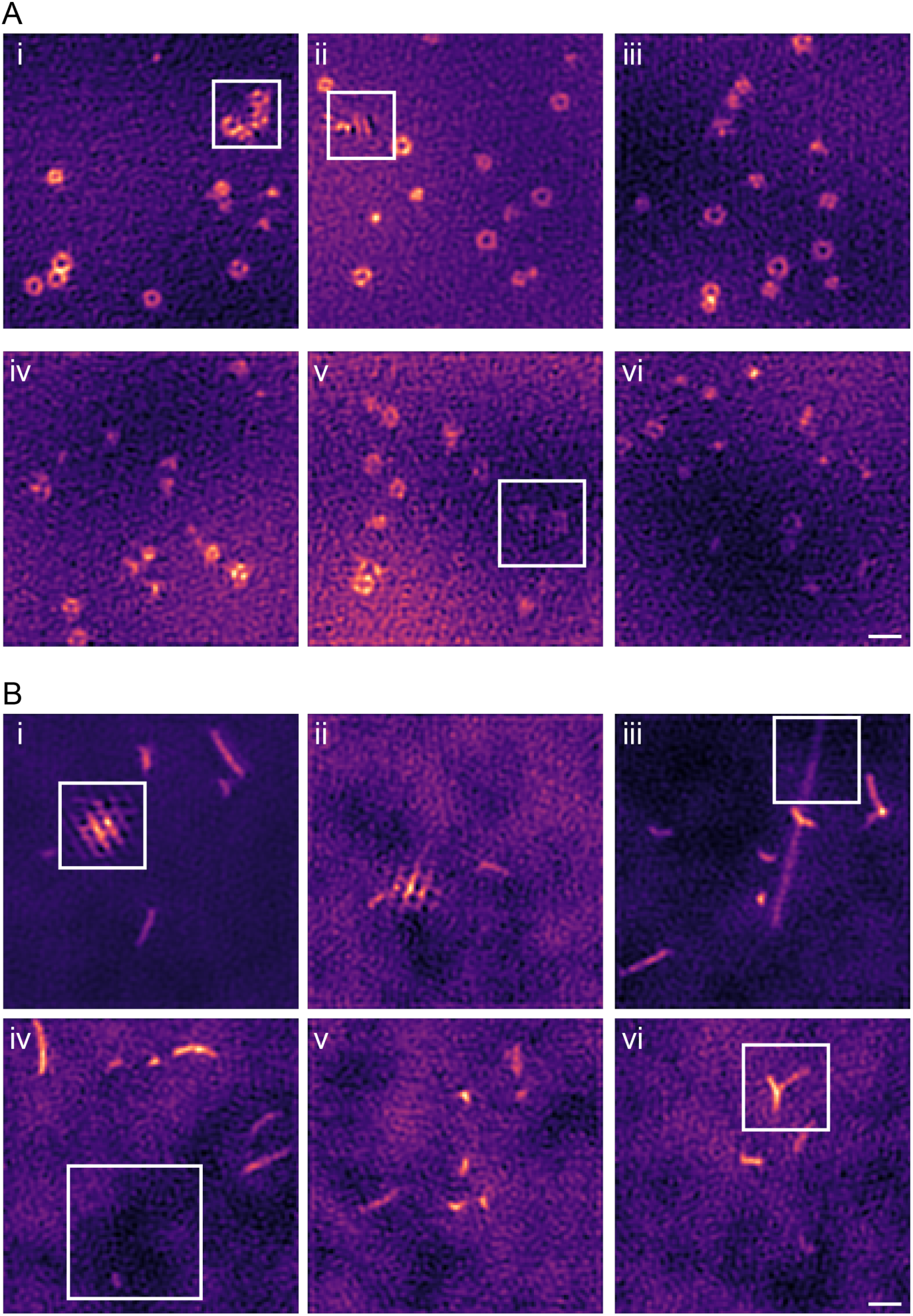
Synthetic TIRF-SIM training data spanning the full range of carrier morphologies, densities, and imaging conditions. Reconstructed synthetic TIRF-SIM image patches (128 × 128 px) generated to train Shape2Fate detectors for endocytic and exocytic carriers. **(A)** Six representative patches *(i–vi)* from the CME training set illustrate annular and disk-like CCPs across a range of densities, clustering, artifacts and SNR levels. **(B)** Six representative patches *(i–vi)* from the exocytosis training set contain mixtures of compact vesicles, elongated tubular carriers and diffusing cargo at the PM. In both panels, white boxes highlight features synthesized to mimic structures and reconstruction artifacts observed in real TIRF-SIM data, using the physics-based image-formation model described in the main text. Examples include partial annuli and clusters **(A *i*)**, motion-blurred objects **(A *ii*)**, static autofluorescent structures **(B *iii*)** and irregular background and membrane attachment **(B *iv*).** These examples illustrate the range of shapes, noise levels and imaging irregularities encompassed by the synthetic training library used to train Shape2Fate detectors. Scale bar: 500 nm.

**Supplementary Figure 2.**
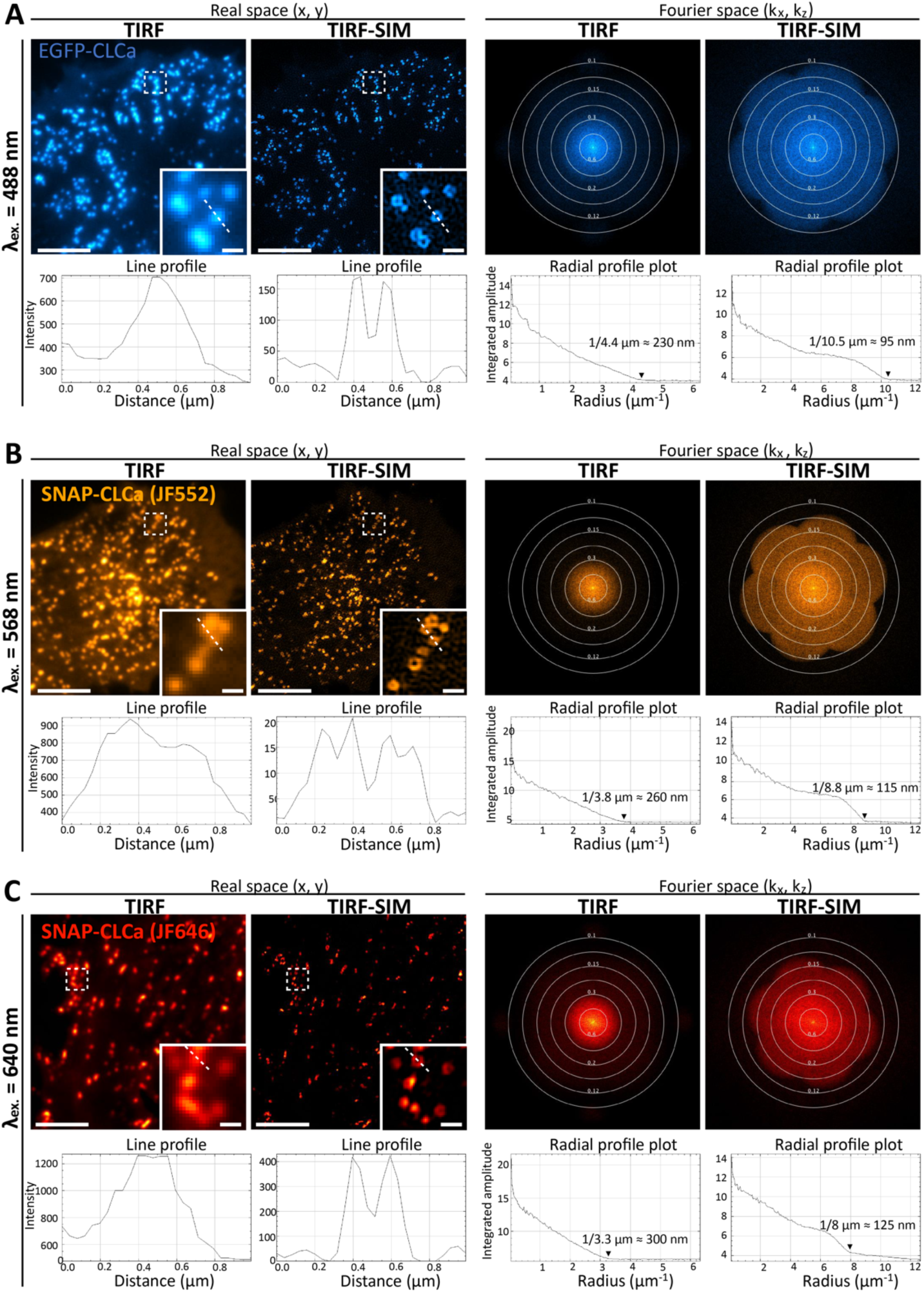
Effective lateral resolution of TIRF-SIM in our experimental conditions on the DeltaVision OMX SR across three excitation wavelengths. Comparison based on real-world data of CCPs labelled with EGFP-CLCa **(A)**, JFX552-conjugated ligand **(B)**, or JFX646-conjugated ligand **(C)**. Representative examples shown with conventional TIRF resolution (obtained from averaging the 9 raw images of a selected frame) and the corresponding reconstructed TIRF-SIM image displayed in real space (left column) and frequency space (right column). Insets show a 5-times magnified view of the boxed region. Scale bars: 10 and 1 µm, respectively. Intensity profile plots of the dotted lines in the inset magnifications highlight the ability of TIRF-SIM to resolve the annular cross-section and internal lumen of CCPs in contrast to conventional TIRF. Radial frequency profile of the Fourier amplitude plots highlights the increased frequency range of TIRF-SIM with arrows marking the inflection point at the estimated practical resolution cutoff.

**Supplementary Figure 3.**
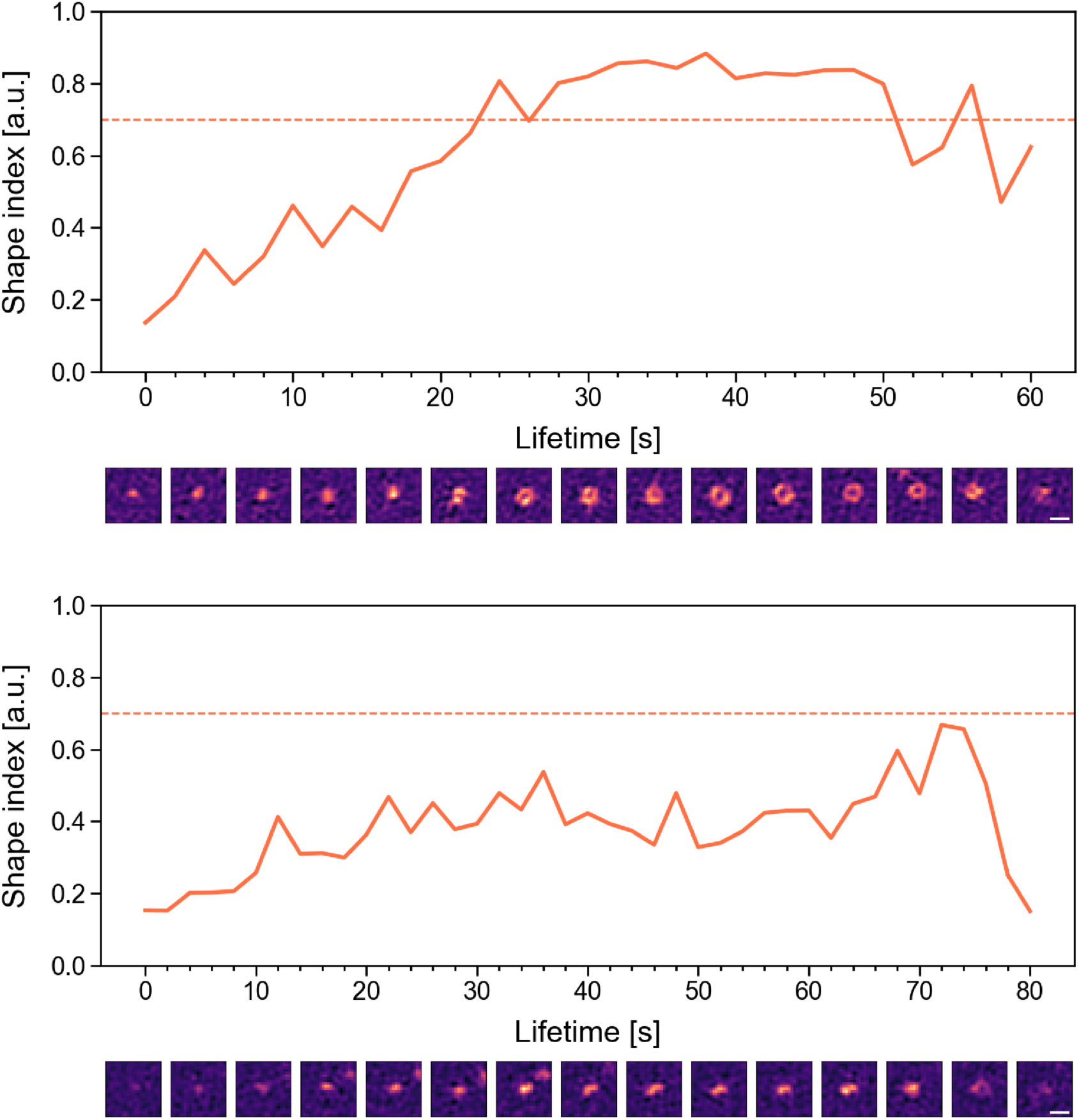
Example of Shape2Fate productivity classification illustrating Shape Index trajectory evolution for a canonical productive CCP and a long-lived non-productive assembly. SI trajectories (orange) and corresponding TIRF-SIM cutouts for a productive (top) and an abortive (bottom) CCP. Dashed orange line marks the productivity threshold (SI = 0.7). In the productive example, the SI rises from near zero (for a nascent assembly unresolved at TIRF-SIM resolution) through the threshold as an annular morphology develops. The SI then drops sharply upon vesicle departure from the evanescent field following scission. The abortive example illustrates a long-lived clathrin assembly, likely a flat or stalled coat (see ref. [5, 8]), whose SI fluctuates at low-to-intermediate values without reaching the productivity threshold. Despite its extended lifetime, which could confound intensity- or lifetime-based classifiers, Shape2Fate correctly assigns this event as abortive on the basis of its incomplete SI progression. Scale bar: 250 nm.

**Supplementary Figure 4.**
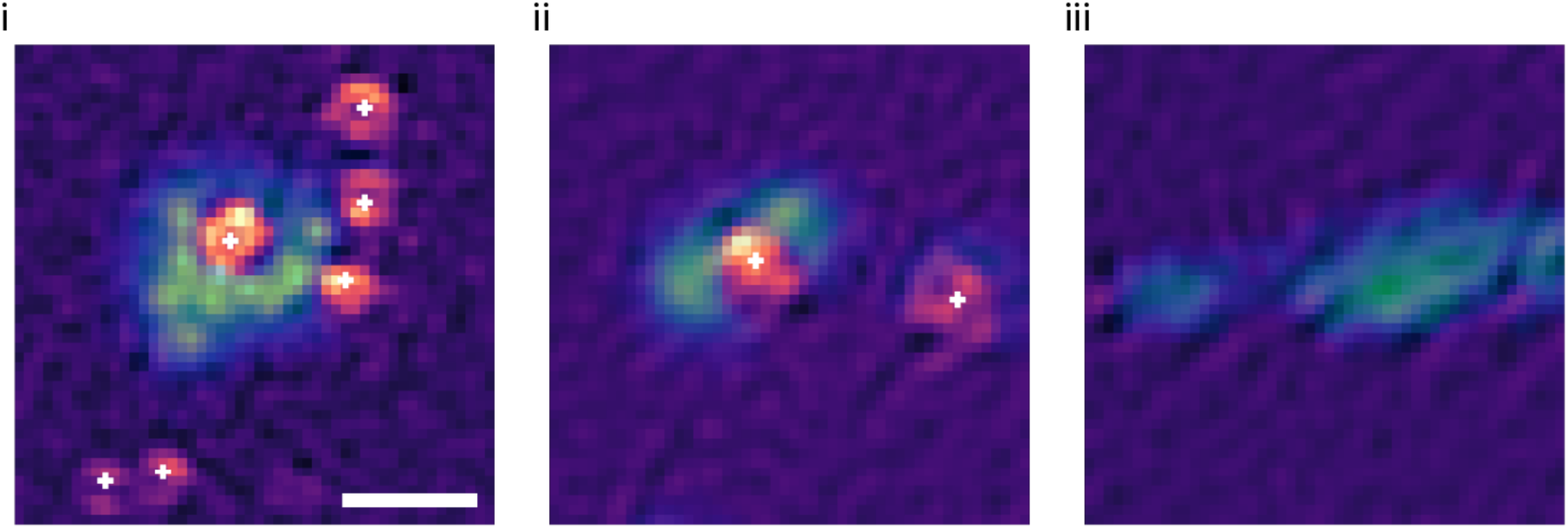
Residual clathrin signal in TIRF-SIM not accounted for by Shape2Fate CCP detections. White crosses mark successfully detected CCPs in combination with unresolved signal visualized in a blue/green hue: *(i)* a dense cluster in which individual CCPs are resolved but overlapping signal remains unassigned; *(ii)* a non-canonical clathrin structure from which only CCP-like sub-regions are detected; and *(iii)* an elongated signal lacking any detections, likely a SIM reconstruction artifact from object motion during multi-frame acquisition. Scale bar: 500 nm

**Supplementary Figure 5.**
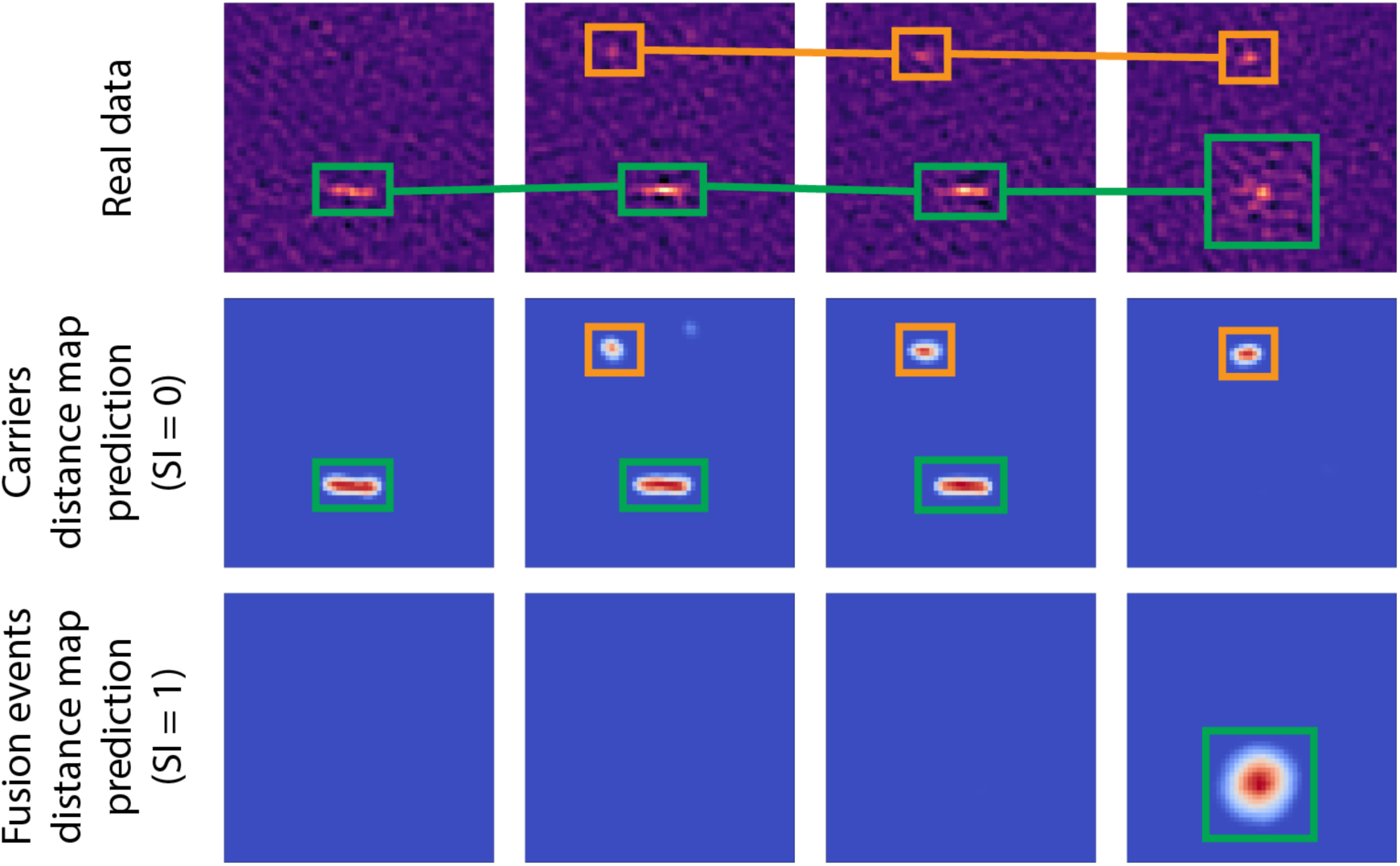
Shape2Fate exocytosis pipeline applied to real TIRF-SIM data: detection, linking, and preliminary productivity classification. The trained U-Net detector is applied to real TIRF-SIM frames of exocytic carriers (top row), predicting two distance maps per frame: one for intact carriers (SI = 0; middle row) and one for fusion events (SI = 1; bottom row). Detections are extracted from each map by connected component analysis and linked across frames into trajectories (coloured boxes and connecting lines in top row). Two example trajectories illustrate distinct carrier fates. The orange-boxed carrier is detected across all frames in the carrier channel but produces no fusion signal, yielding a non-productive trajectory. The green-boxed carrier is similarly tracked in the carrier channel until the final frame, where an isotropic signal burst appears in the fusion event map, consistent with cargo release. This trajectory is flagged as potentially productive and passed to the auxiliary fusion classifier for verification on the raw low-resolution TIRF-SIM frames (see Methods).

**Supplementary Figure 6.**
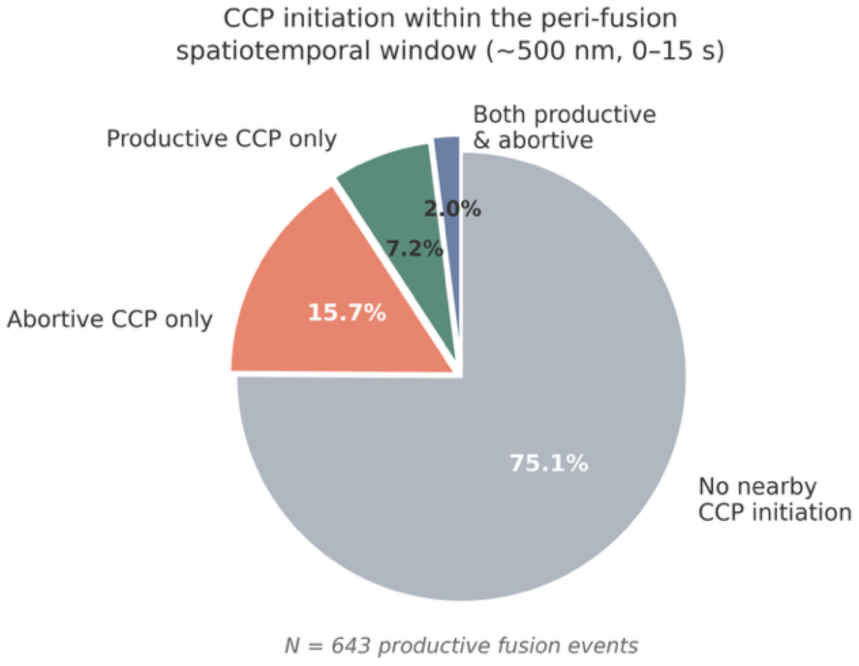
Fraction of productive exocytic fusion events followed by CCP initiation during synchronized constitutive secretion in SH-SY5Y cells using the RUSH system. Pie chart classifying each productive fusion event in SH-SY5Y cells by whether a CCP initiated within the surrounding spatiotemporal window (see Fig. 6A, B for window definition): a productive CCP only (7.2%, green), an abortive CCP only (15.7%, orange), both productive and abortive CCPs (2.0%, hatched), or no CCP (75.1%, grey). Approximately one quarter of fusion events triggered at least one nearby CCP initiation. N = 643 productive fusion events.

**Supplementary Figure 7.**
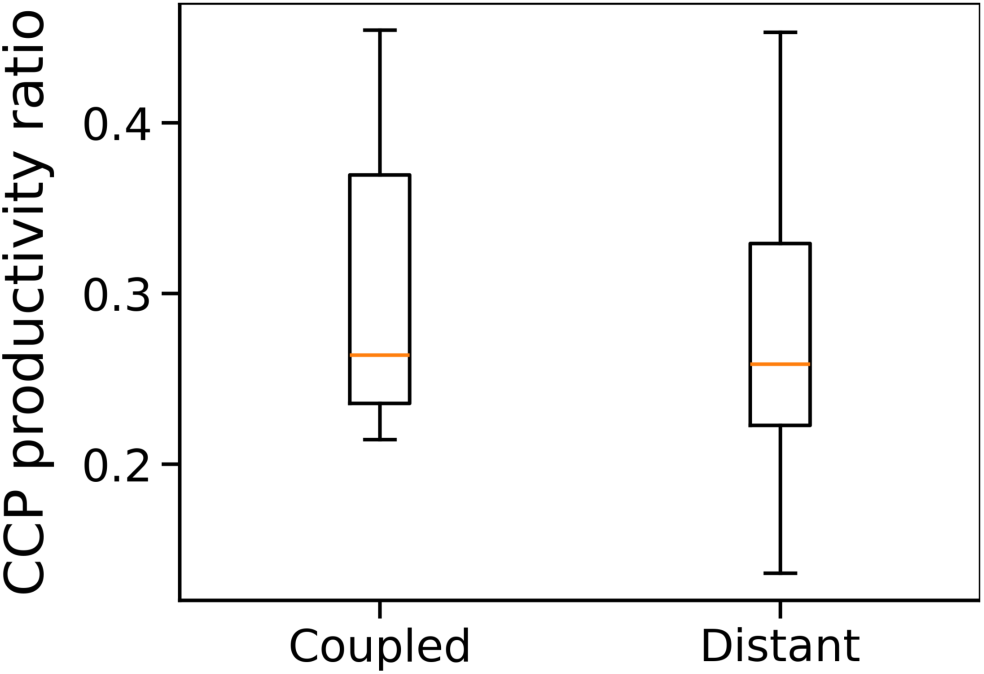
Maturation efficiency of CCPs initiated near productive exocytic fusion sites versus elsewhere in SH-SY5Y cells using the synchronized RUSH system. Box plots comparing the CCP productivity ratio (productive/total trajectories) of coupled CCPs, defined as those initiated within the spatiotemporal window surrounding productive fusion events (see Fig. 6A), with that of distant CCPs in SH-SY5Y cells expressing mEmerald–LAMP1 and mRuby2–CLCa. No significant difference was observed (Welch’s t-test, p = 0.57), indicating that although more CCPs initiate near fusion sites (Fig. 6), their probability of completing productive endocytosis is constant.

**Supplementary Figure 8.**
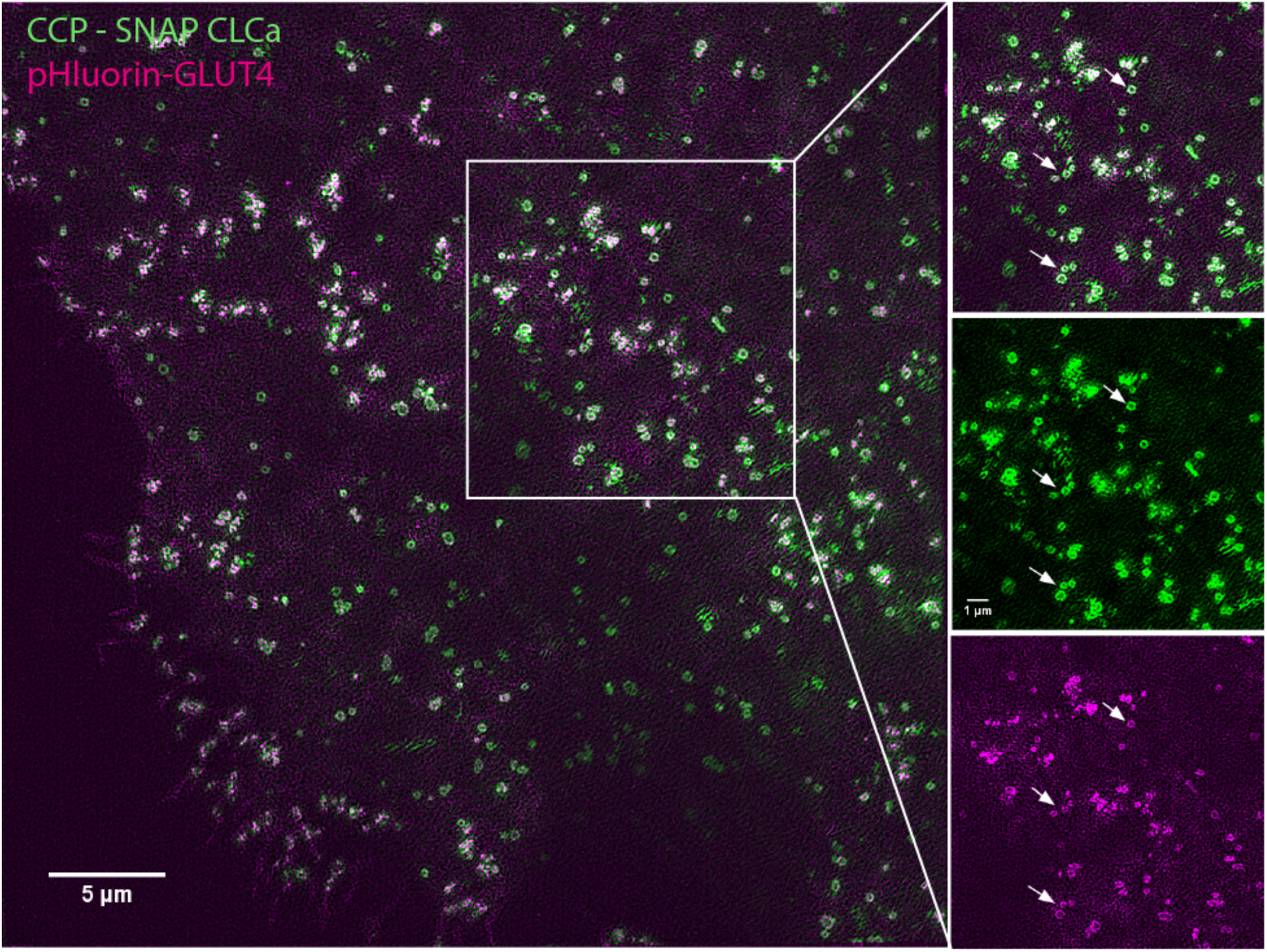
Spatial co-localization of newly delivered GLUT4 with pre-existing CCPs in an insulin-stimulated adipocyte. Dual-channel TIRF-SIM snapshot of a 3T3-L1 adipocyte 5 minutes after insulin stimulation, representative of the time-lapse shown in Supplementary Movie 5. Green: SNAP–CLCa, marking CCPs; magenta: pHluorin–GLUT4, marking transporter inserted at the PM by GSV fusion. Co-localization of GLUT4 signal with CCPs appears as white overlap. Left: whole-cell overview; boxed region is magnified on the right as an overlay (top), the CLCa channel alone (middle), and the GLUT4 channel alone (bottom). Arrows indicate individual representative sites where pHluorin–GLUT4 fluorescence accumulates at or immediately adjacent to CCPs, consistent with the rapid post-fusion cargo capture quantified in Fig. 7G.

**Supplementary Figure 9.**
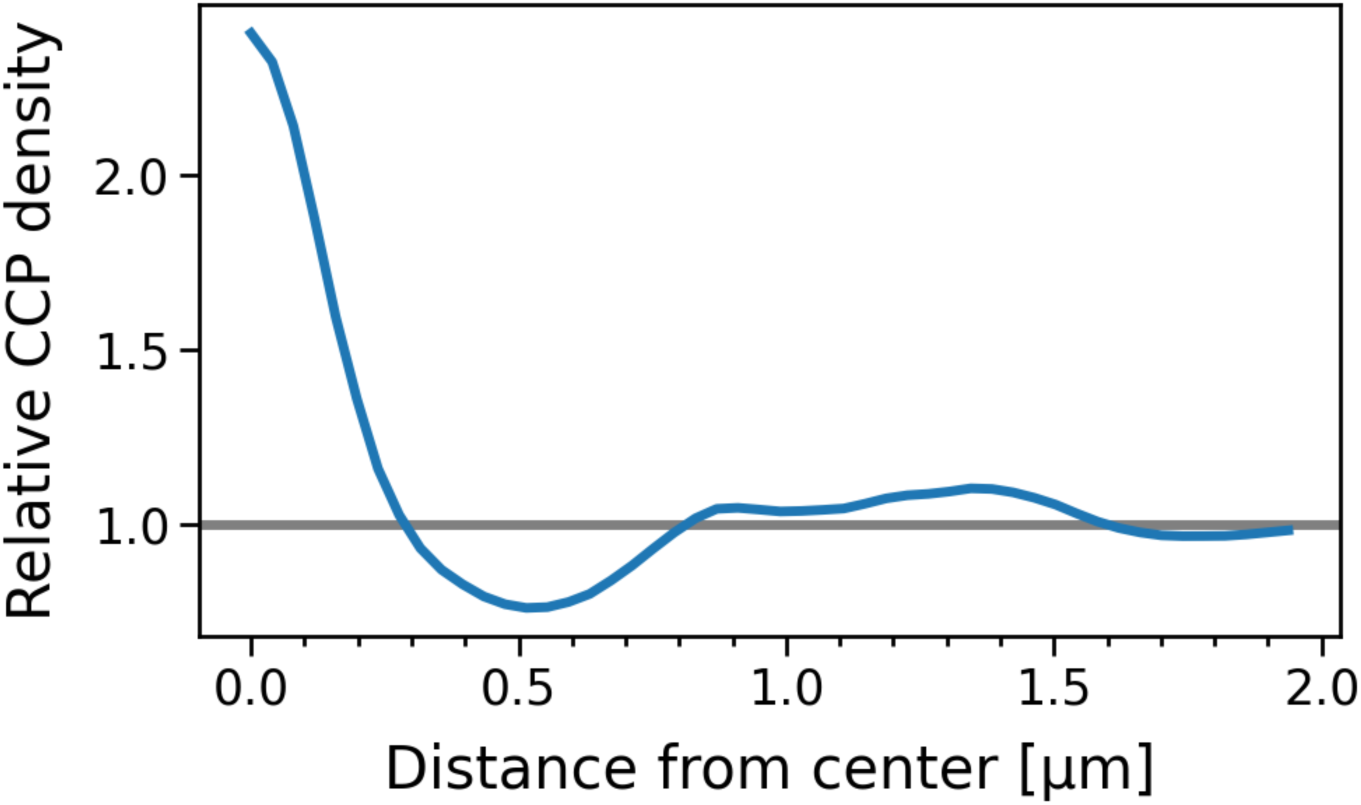
Radial profile of CCP density around GSV fusion sites in insulin-stimulated adipocytes. One-dimensional radial average of the fusion-centred CCP density heatmap shown in Fig. 7D, plotted as relative CCP density (normalized to the cell-wide average; grey line = 1) versus distance from the GSV fusion centre. CCP density is elevated near fusion sites and approaches baseline at larger distances, consistent with a spatially confined CCP-enriched zone surrounding productive exocytic events.

**Supplementary Figure 10.**
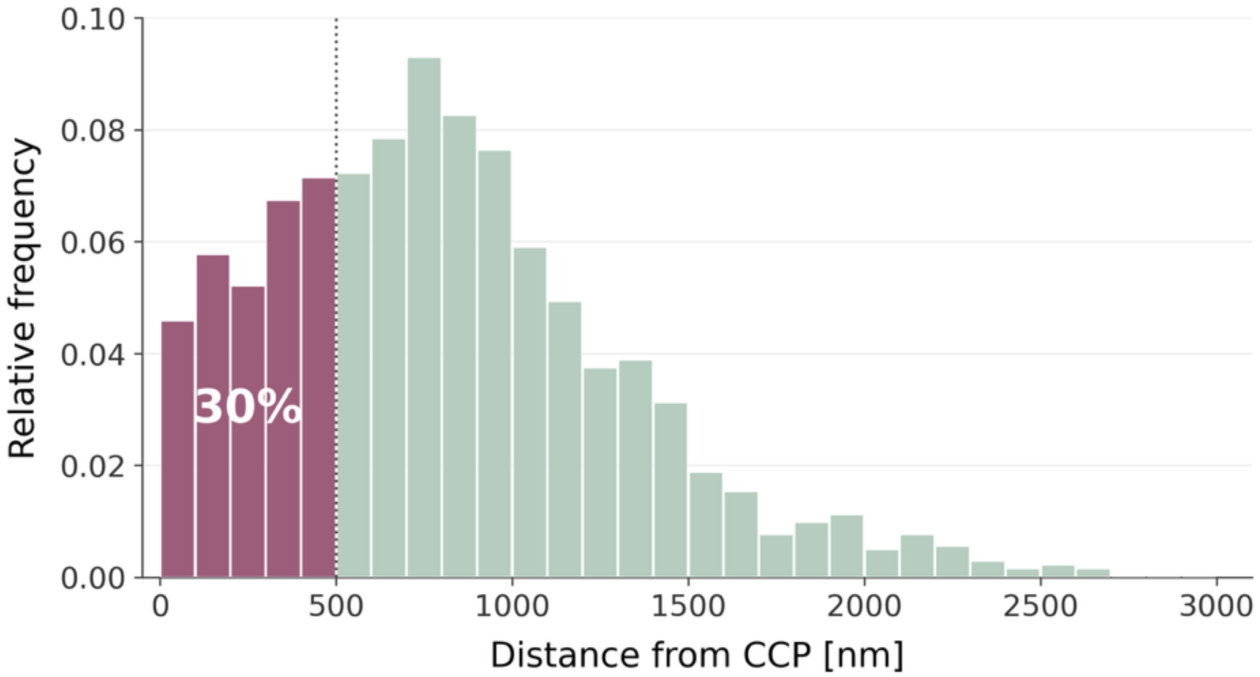
Distances of productive GLUT4 exocytic fusion events to nearest CCP. About 30% of fusions occur within 500 nm of a CCP (green).

**Supplementary Figure 11.**
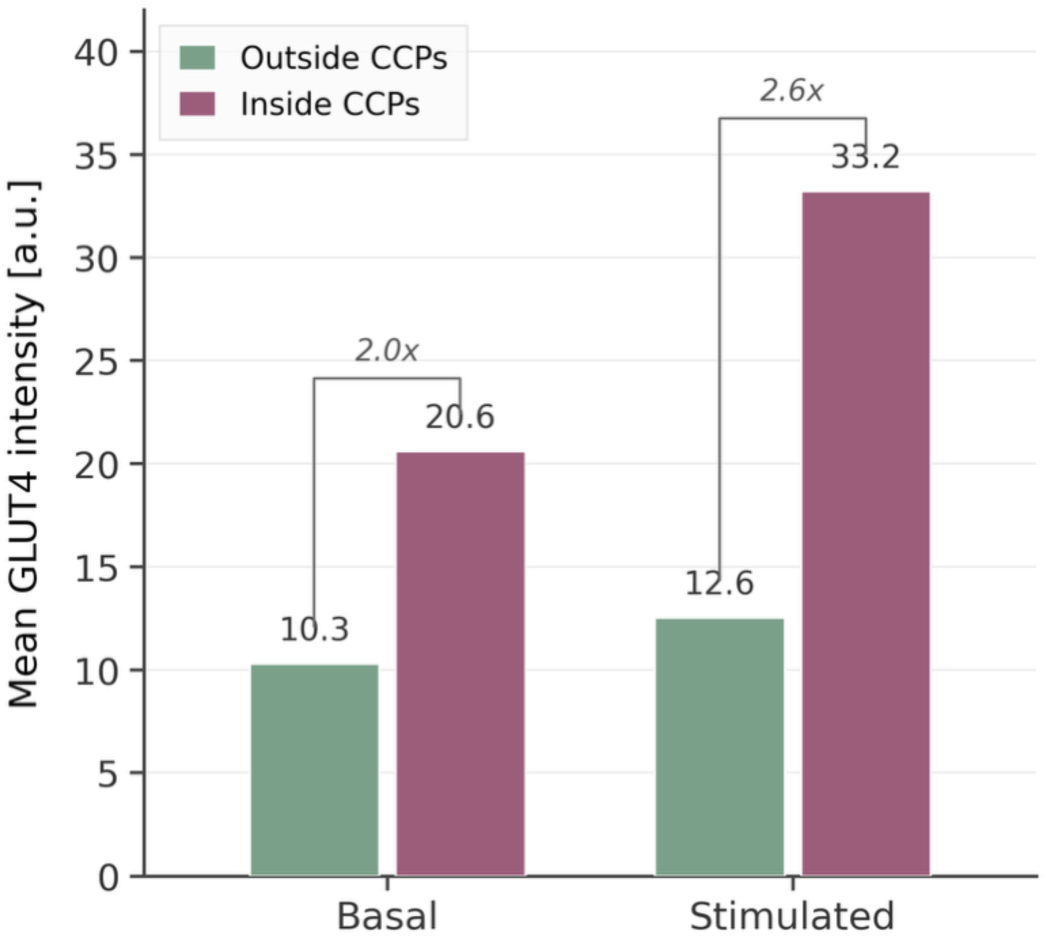
GLUT4 is preferentially enriched at CCPs, and this enrichment increases upon insulin stimulation. Mean GLUT4 intensity (pHluorin–GLUT4, arbitrary units) measured at PM regions inside CCPs (plum) versus outside CCPs (sage) under basal and insulin-stimulated conditions. GLUT4 intensity inside CCPs is approximately two-fold higher than outside under basal conditions (2.0×) and increases to 2.6× after insulin stimulation, consistent with the targeted delivery of GSVs to pre-existing CCP sites resolved by Shape2Fate (Fig. 7C–G). Data pooled from n = 3 independent replicates.

**Supplementary Figure 12.**
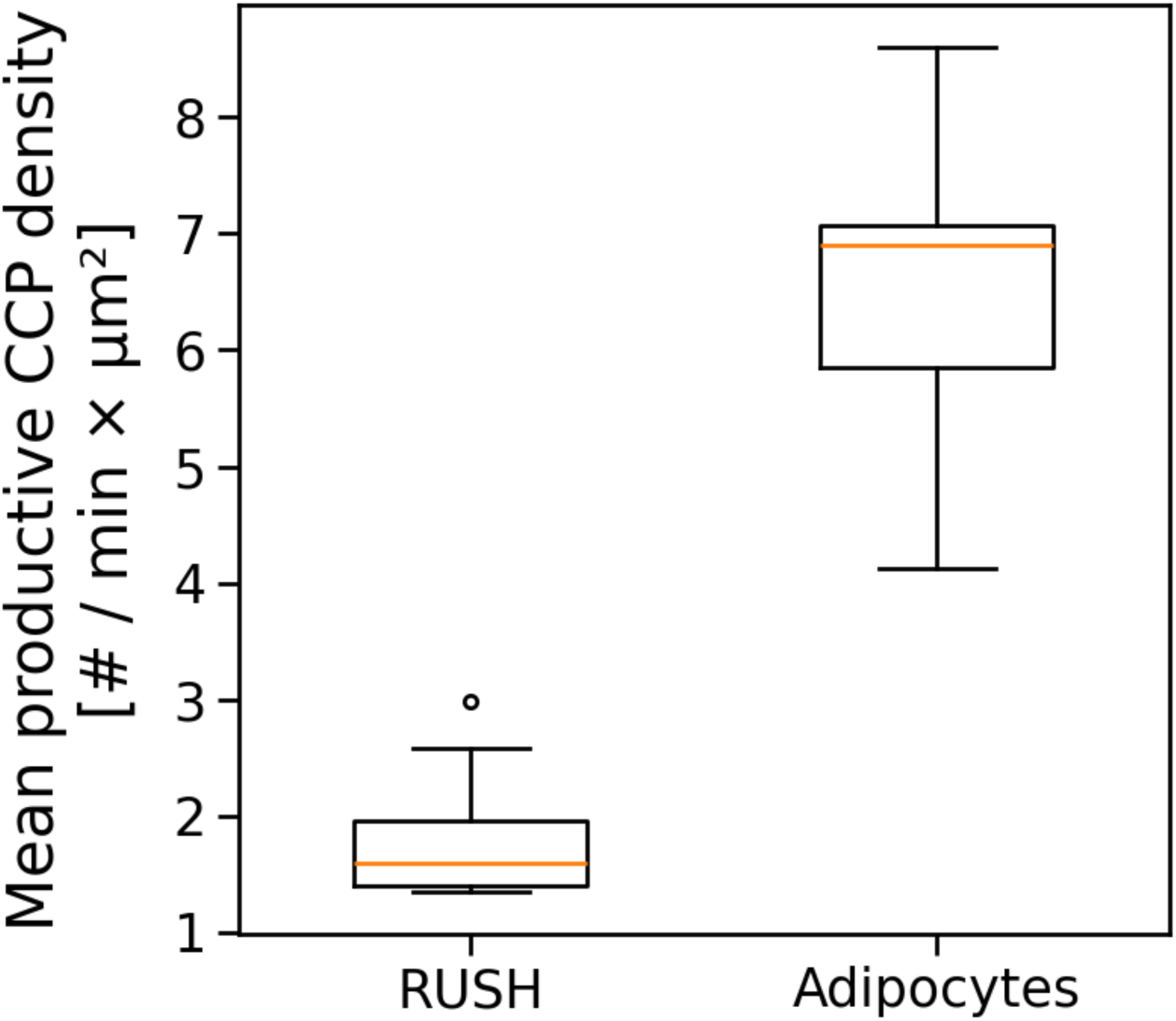
Productive CCP density during exocytosis in SH-SY5Y RUSH cells and insulin-stimulated adipocytes. Boxplots show the mean density of productive CCPs per area and time. Adipocytes exhibit significantly higher densities compared to RUSH cells (p = 0.00005).

**Supplementary Table 1.** Quantitative summary of CME tracking performance on validation dataset. Values are reported as the mean across all annotators.

|  | Annotators | Shape2Fate | Shape2Fate<br>(without SI) | cmeAnalysis | TraCKer |
| --- | --- | --- | --- | --- | --- |
| MOTA | 0.77 | 0.79 | 0.79 | 0.50 | 0.59 |
| 1 – MOTP | 0.68 | 0.64 | 0.64 | 0.40 | 0.33 |
| HOTA | 0.79 | 0.77 | 0.76 | 0.58 | 0.60 |
| DetA | 0.81 | 0.82 | 0.82 | 0.58 | 0.64 |
| AssA | 0.77 | 0.72 | 0.70 | 0.58 | 0.56 |
| μTIOU | 0.65 | 0.64 | 0.62 | 0.40 | 0.40 |

**Supplementary Table 2.** Quantitative summary of CME tracking performance on test dataset. Tracking performance was evaluated at a representative frame rate of ∼2 s. Reported values are averaged across all possible frame subsets and shown as mean ± std.

|  | Shape2Fate | cmeAnalysis | TraCKer |
| --- | --- | --- | --- |
| MOTA | 0.75 ± 0.007 | -0.06 ± 0.016 | 0.63 ± 0.006 |
| 1 – MOTP | 0.60 ± 0.005 | 0.54 ± 0.004 | 0.40 ± 0.007 |
| HOTA | 0.72 ± 0.007 | 0.45 ± 0.006 | 0.64 ± 0.010 |
| DetA | 0.79 ± 0.006 | 0.42 ± 0.005 | 0.68 ± 0.006 |
| AssA | 0.65 ± 0.014 | 0.47 ± 0.011 | 0.60 ± 0.020 |
| μTIOU | 0.61 ± 0.012 | 0.29 ± 0.007 | 0.53 ± 0.008 |

**Supplementary Table 3.**
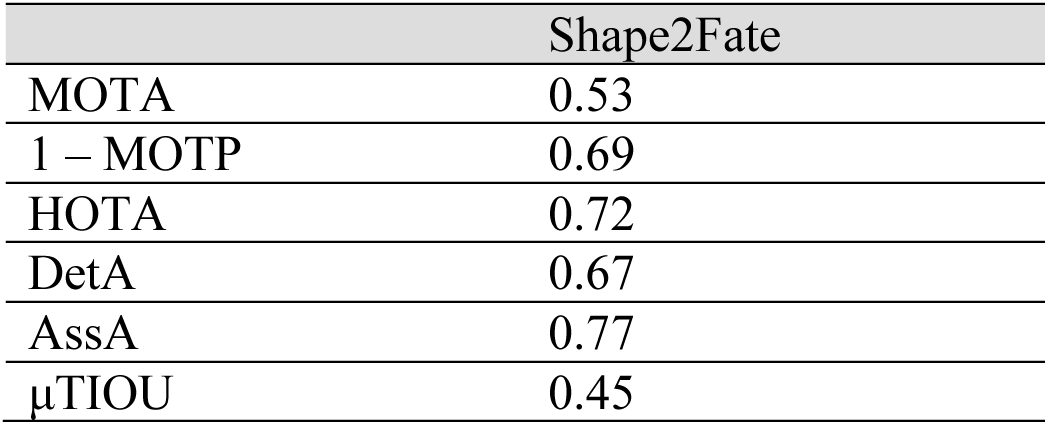
Quantitative summary of exocytosis tracking performance.

|  | Shape2Fate |
| --- | --- |
| MOTA | 0.53 |
| 1 – MOTP | 0.69 |
| HOTA | 0.72 |
| DetA | 0.67 |
| AssA | 0.77 |
| μTIOU | 0.45 |

**Supplementary Table 4.** Quantitative summary of productive exocytosis classification. To ensure fair comparison, different strategies for suppressing structured-illumination interference in raw low-resolution TIRF-SIM frames were tested: using raw frames (raw), averaging three phase-offset frames of the same orientation (avg 3), or averaging nine frames in a sliding window shifted by one frame (slid. win.).

|  | Shape2Fate | IVEA (raw) | IVEA (avg 3) | IVEA (slid. win.) |
| --- | --- | --- | --- | --- |
| Accuracy | 0.93 | 0.68 | 0.74 | 0.67 |
| Precision | 0.89 | 0.15 | 0.26 | 0.03 |
| Recall | 0.90 | 0.59 | 0.87 | 0.89 |
| F1 | 0.90 | 0.23 | 0.40 | 0.06 |

**Supplementary Movie 1. Qualitative example of tracking performance of automated methods and human annotators.**

Comparison of CCP trajectories generated by Shape2Fate, other automated trackers, and human annotations on representative TIRF-SIM or TIRF data. Similar trajectories are color-matched across panels. Scale bar: 500 nm.

**Supplementary Movie 2. Qualitative example of endocytosis and exocytosis tracking.** Representative CME (top row, EGFP-CLCa) and RUSH exocytosis (bottom row, mEmerald-LAMP1) sequences from different cells are shown. Raw movies are displayed on the left, and the same sequences overlaid with trajectories generated by Shape2Fate are shown on the right. Trajectories are color-coded to differentiate between productive (yellow) and non-productive (cyan) events. Both sequences were recorded at different frame rates and are not aligned. Scale bar: 1 µm.

**Supplementary Movie 3. Qualitative examples of endo-exo coupling in RUSH cells.** Representative sequences from SH-SY5Y cells showing spatial and temporal coupling of exocytic carrier fusion with nearby CCP initiation. Two channels are displayed: exocytosis (magenta, mEmerald-reporter, 400 ms/frame) and CME (green, SNAP-CLCa, 3.2 s/frame). The CME channel was upsampled to match the exocytic frame rate by linear interpolation. A timer indicates the moment of fusion (t = 0), and key phases of the coupling process are highlighted. In the top-left quadrant, the corresponding plot of SI over time is shown for both endocytic and exocytic events. Image contrast was artificially enhanced for visibility. Scale bar: 500 nm.

**Supplementary Movie 4. Confirmation of successful adipocyte differentiation and insulin responsiveness by pHluorin-GLUT4 translocation.** TIRF time-lapse acquisition (20 min, 30 s per frame) of pHluorin-GLUT4 in day-nine differentiated 3T3-L1 adipocytes following stimulation with 100 nM insulin. The rapid, cell-wide accumulation of pHluorin-GLUT4 signal at the PM immediately after insulin addition serves as a functional control for successful adipocyte differentiation, as insulin-responsive GSV translocation to the PM is a hallmark of the mature adipocyte phenotype and is largely absent in undifferentiated fibroblasts. This confirms that the differentiated adipocytes used for TIRF-SIM exo–endocytic coupling experiments (Fig. 7, Supplementary Movies 5–6) are insulin-responsive and competent for regulated GLUT4 exocytosis. Scale bar: 2 µm.

**Supplementary Movie 5. Whole-cell TIRF-SIM imaging of insulin-stimulated GLUT4 exocytosis and CME in an adipocyte.** Representative dual-colour TIRF-SIM movie of a single adipocyte 5 minutes following insulin stimulation. The CME channel was upsampled to match the exocytic frame rate by linear interpolation. Magenta: GSVs (labelled with pHluorin–GLUT4), visualising fusion events and subsequent lateral diffusion of newly inserted transporter at the PM. Yellow: SNAP-CLCa visualizing CCPs. The movie, 3 min long acquired with interleaved acquisition at a frame rate 250 ms/frame (pHluorin–GLUT4) and at a frame rate 2 s/frame with SNAP-CLCa, illustrates the spatial proximity of productive GLUT4 fusion events to pre-existing CCPs and the rapid accumulation of newly delivered GLUT4 within CCP following fusion, consistent with a coupling architecture in which exocytic carriers target pre-formed endocytic sites for immediate cargo capture.

**Supplementary Movie 6. Qualitative examples of endo-exo coupling in insulin-stimulated adipocytes.** Representative sequences from 3T3-L1 adipocytes showing coordinated exocytic fusion near pre-formed CCPs under insulin stimulation. Two channels are displayed: exocytosis (magenta, pHluorin-GLUT4, 250 ms/frame) and CME (green, SNAP-CLCa, 2 s/frame). The CME channel was upsampled to match the exocytic frame rate by linear interpolation. A timer indicates the moment of fusion (t = 0), and key phases of the coupling process are highlighted. Image contrast was artificially enhanced for visibility. Scale bar: 500 nm.

